# Perceptual sensitivity is explained by optimization for ecological tasks

**DOI:** 10.64898/2026.09.28.755158

**Authors:** Mark R. Saddler, Torsten Dau, Josh H. McDermott

## Abstract

When asked to discriminate between two stimuli that differ along some dimension, such as intensity, frequency, or orientation, humans exhibit a smallest stimulus difference, or threshold, below which discrimination becomes impossible. These limits on perceptual sensitivity have traditionally been explained by internal noise. Contemporary theories instead link perceptual sensitivity to efficient coding of the natural environment but have not been comprehensively tested. Here, we measured the psychoacoustic sensitivity of a computational model optimized for everyday sound localization and recognition tasks. We used linear classifiers operating on the model’s learned representations to estimate 1632 thresholds spanning the perceptual dimensions of sound. These model-derived thresholds closely matched thresholds measured behaviorally in humans on the same test battery. The results show that many classical psychophysical phenomena emerge from representations optimized for natural behavior and suggest that discrimination thresholds are determined by linear separability in these optimized representations, rather than intrinsic neural noise.

## INTRODUCTION

Our perceptual systems are generally adept at sensing and interpreting the world around us but nonetheless have limits to their acuity. For more than a century, psychophysical thresholds – measurements of the smallest stimulus differences humans can perceive – have been the primary means of characterizing these limits.

Thresholds place strong constraints on theories of sensory processing. Differences in perceptual sensitivity along stimulus dimensions, such as spatial frequency^1^ and color^2^ in vision, and interaural phase^3^ and temporal modulation^4^ in audition, have long been interpreted as evidence for specialized neural mechanisms. However, less attention has been given to the absolute magnitude of perceptual sensitivity: why are organisms not more or less sensitive to stimulus differences?

Traditional mechanistic models often explain perceptual limits by positing internal noise^3,5,6^. To quantitatively predict human thresholds, these models typically assume sources of variability motivated by the stochastic nature of biological hardware, including variability in neural spiking, synaptic transmission, and molecular processes^7^. Under this view, perceptual sensitivity is fundamentally constrained by the unreliability of neural machinery.

An alternative possibility is that perceptual limits reflect what is needed to enable natural behavior^8^. Efficient coding theories propose that sensory systems allocate limited biological resources according to environmental statistics while reducing redundancy in sensory representations^8–13^. Commonly occurring stimuli may thus be encoded with higher fidelity. More recent instantiations of efficient coding theories further argue that efficient codes determine the best achievable discrimination thresholds^14,15^. However, efficient coding theories are not presently able to derive representations of multiple stimulus dimensions, making the full implications of such theories difficult to test.

By contrast, deep learning provides a way to derive rich multi-dimensional representations that subserve tasks and has enabled a new generation of perceptual models. Models optimized for real-world sensory tasks capture many aspects of human perceptual behavior, including speech recognition^16,17^, object recognition^18,19^, pitch perception^20^, sound localization^21^, selective auditory attention^22^, and perceptual confidence^23^. Here we ask whether task-optimized representations can account for the quantitative limits of human perception.

A priori it seemed possible that natural behaviors might substantially constrain the precision with which sensory information is represented. To support such behaviors, sensory systems must preserve stimulus differences that are behaviorally relevant, while also having little incentive to represent differences smaller than those that improve the performance of ecological tasks. Consistent with this idea, task optimization via gradient descent has been argued to create representations that embody efficient coding by being more sensitive to commonly occurring stimulus features, and to accordingly determine the discriminability of basic features^24^. However, previous tests of this proposal have been limited to qualitative tests for three visual dimensions (orientation, hue, and spatial frequency)^24^, leaving it unclear whether task optimization can broadly account for perceptual sensitivity.

Here, we present a model optimized solely for naturalistic auditory behavior and test whether such optimization is sufficient to produce the qualitative and quantitative characteristics of human perceptual sensitivity. The model was jointly trained to localize and recognize words, voices, and environmental sounds from simulated auditory nerve representations of natural scenes. We then measured 1632 psychoacoustic thresholds from the model across 31 experiments, assessing sensitivity along the principal dimensions of auditory perception as well as their interactions. The results suggest the characteristic limits of human hearing can largely be explained by perceptual representations optimized for everyday listening behavior.

## RESULTS

### Model construction

To obtain a model capable of performing everyday hearing tasks, we trained a deep neural network to jointly localize and recognize words, voices, and environmental sounds from simulated auditory nerve representations of naturalistic scenes (Fig. 1a). These tasks were chosen because they represent common listening behaviors that plausibly shape auditory perception and can readily be operationalized with supervised machine learning^17,25^. We do not claim these tasks are uniquely important; rather, they provide a representative sample of real-world auditory behavior. The network consisted of a shared trunk of six convolutional layers followed by four parallel task-specific branches, each consisting of four convolutional layers. This architecture encouraged the development of representations useful across multiple behaviors while allowing specialization for individual tasks^16^. All model parameters were optimized solely for performance on these ecological tasks, and the model was never fit to human behavioral data.

**Figure 1.**
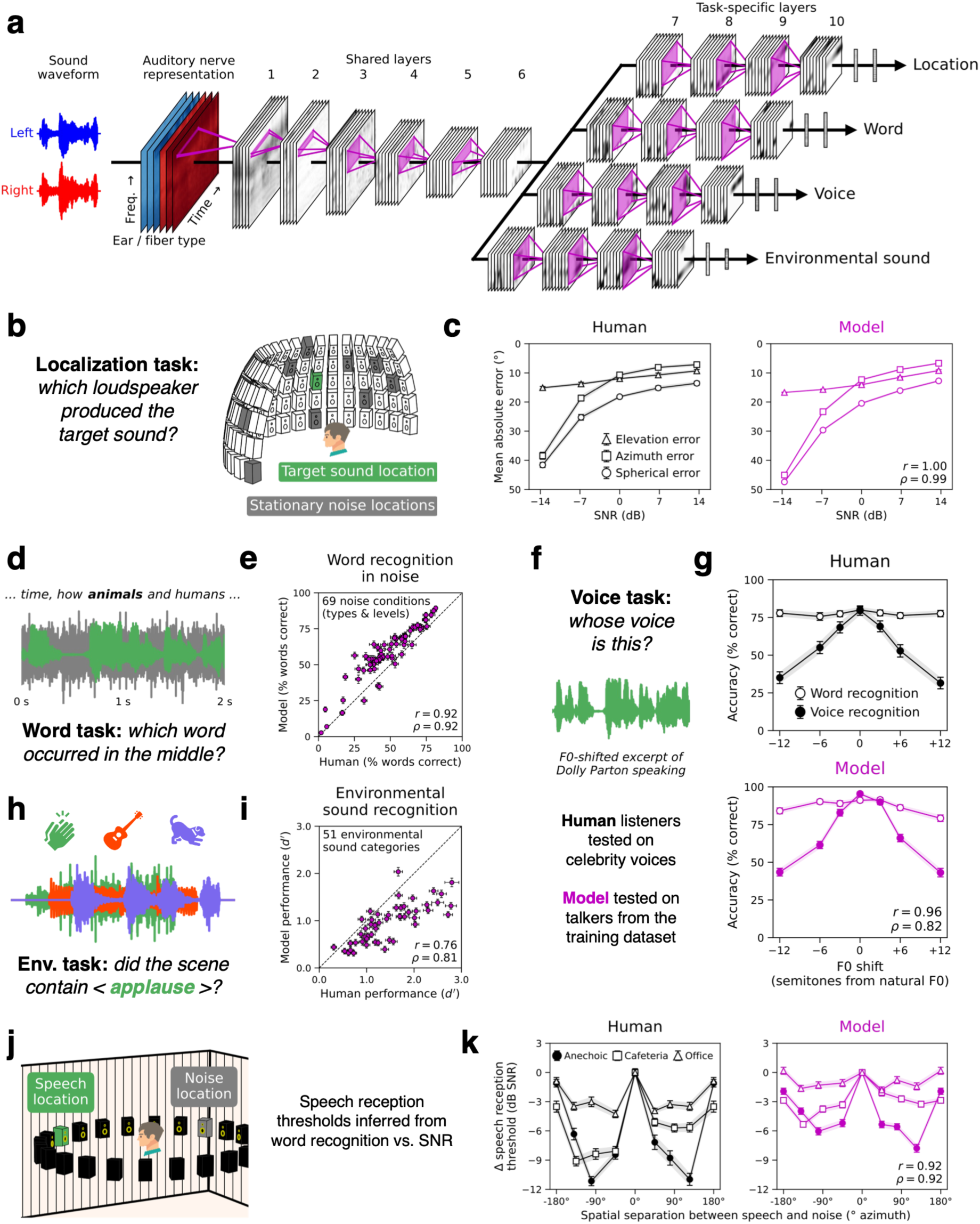
A model optimized for ecological hearing tasks reproduces characteristics of human sound localization and recognition behavior. **(a)** Schematic of the convolutional neural network trained to localize and recognize words, voices, and environmental sounds from simulated auditory nerve representations. **(b)** Sound localization in noise task. **(c)** Comparison of human and model sound localization performance, assessed by measuring mean absolute azimuth, elevation, and spherical error as a function of signal-to-noise ratio. **(d)** Word recognition in noise task. **(e)** Comparison of human and model word recognition accuracy across 69 background noise conditions. **(f)** Familiar voice recognition task. **(g)** Human and model word and voice recognition accuracy measured using speech from familiar talkers whose voice has been pitch-shifted by increasing or decreasing fundamental frequency (F0). **(h)** Environmental sound recognition task. **(i)** Comparison of human and model environmental sound recognition performance across 51 different sound classes. **(j)** Speech reception thresholds were measured for different spatial configurations of speech and noise sources. **(k)** Human and model speech reception thresholds measured as a function of the spatial separation between speech and noise in three acoustically distinct rooms. Human-model similarity for each task is quantified using Pearson (r) and Spearman (ρ) correlation coefficients. Human data in (**c**), (**e**), and (**g**) are from ref.^17^. Human data in (**i**) and (**k**) are from refs.^25,27^, respectively. Error bars indicate ±1 s.e.m.

### Model optimized for ecological tasks exhibits human-like behavior on its training tasks

We compared the model’s localization and recognition performance with that of human listeners on matched tasks and stimuli (aside from minor implementation differences; see Methods). The model exhibited qualitatively and quantitatively human-like behavior across a range of auditory benchmarks, including natural sound localization in noise^17^ (Figs. 1b and 1c), word recognition in real-world noise^16,17^ (Figs. 1d and 1e), word and voice recognition with pitch-shifted speech^26^ (Figs. 1f and 1g), environmental sound recognition^25^ (Figs. 1h and 1i), and speech reception thresholds across spatial configurations and acoustic environments^27^ (Figs. 1j and 1k). Previous work has accounted for some of these behaviors with separate models optimized for either localization or recognition tasks^17,25^. Here, we find that a single multi-task model can account for human behavior across domains. Moreover, the model’s speech reception thresholds exhibit human-like spatial release from masking^28^, demonstrating that human-like interactions between spatial hearing and speech recognition emerge in a system optimized for naturalistic listening behavior. These results are all consistent with the possibility that the model converged on human-like solutions to its ecological training tasks.

### Psychoacoustic thresholds measured from the model’s learned representations

To ask whether optimization for ecological perceptual tasks is sufficient to explain the limits of human perception, we measured the model’s psychophysical thresholds. Perceptual sensitivity is commonly characterized with experiments in which participants make binary detection or discrimination judgments about pairs of stimuli that differ along a controlled stimulus dimension^6^. We simulated such experiments by training linear classifiers to perform two-interval two-alternative forced-choice (2I-2AFC) tasks using the model’s learned feature representations (Fig. 2). Classifiers were trained to convergence, and their performance was used to construct psychometric functions from which perceptual thresholds were inferred.

**Figure 2.**
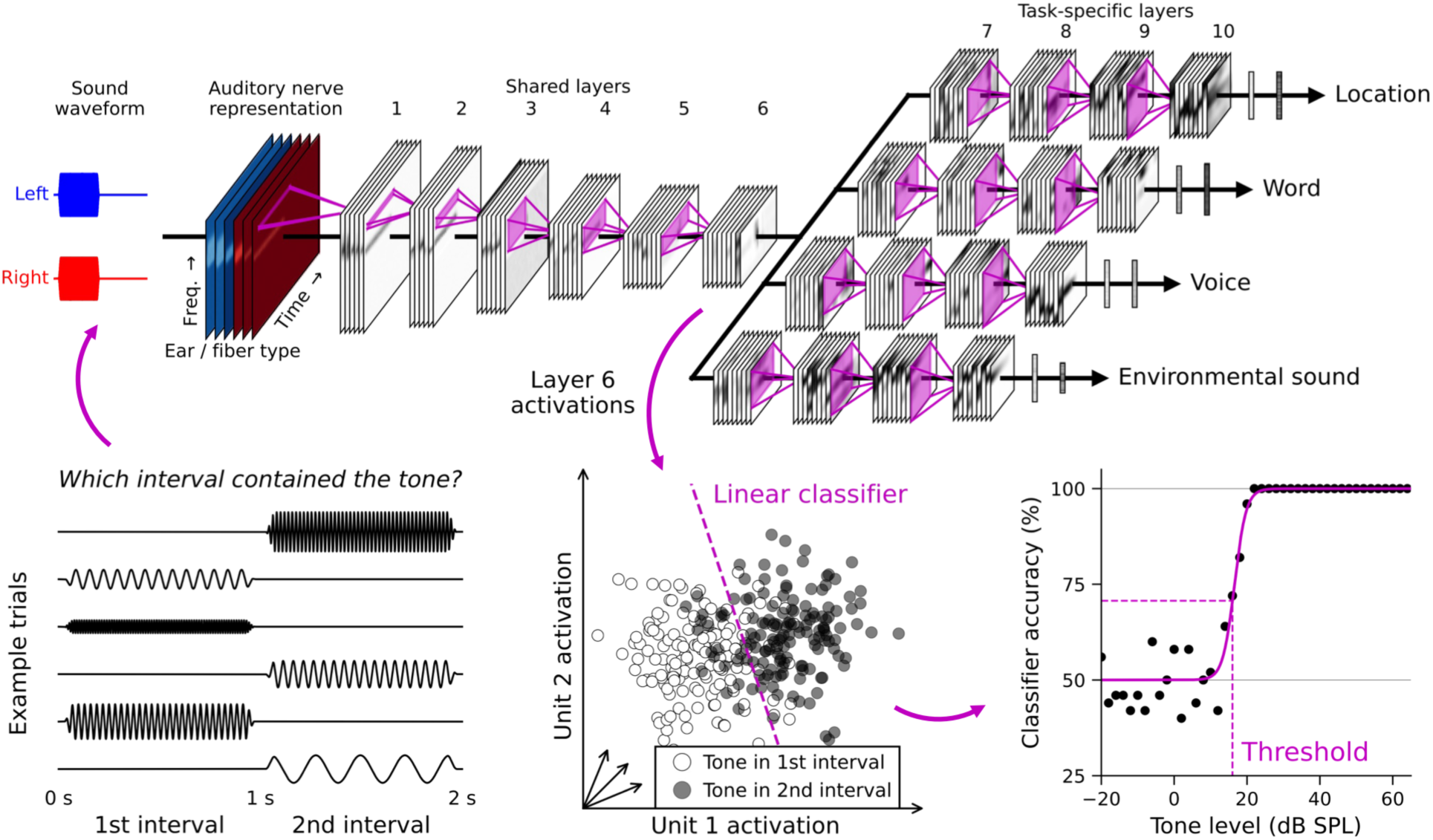
Model psychoacoustic thresholds were estimated via linear classifiers operating on the network’s learned representations. Thresholds were inferred from the performance of binary classifiers trained to perform 2I-2AFC tasks using the model’s layer 6 activations as input.

For instance, to measure audiometric thresholds, we evaluated the model on two-second stimuli, each corresponding to a single two-interval trial. The two intervals were presented in random order, with one containing silence and the other containing a pure tone whose frequency and level varied across trials. We recorded the model’s internal activations for each trial and trained a linear classifier to report whether the tone was in the first or second interval. Thresholds were then estimated from psychometric functions relating classifier performance to tone level at each test frequency. The linear classifier was intended to be analogous to the decision rule a human listener might use to perform the same task using their internal representations. Linear classifiers were chosen on grounds of parsimony, as an unbiased way to evaluate the information made explicit in the model’s representations.

Reasoning that psychoacoustic task judgments are plausibly based on general-purpose auditory representations, we measured thresholds using activations from the deepest model stage shared across the ecological training tasks (layer 6). Results obtained from other model stages are presented at the end of this section.

### Ecologically optimized representations support human-like psychophysical thresholds

We measured 31 different types of thresholds to comprehensively characterize perceptual sensitivity along the four principal dimensions of the auditory input -- intensity, time, frequency, and interaural disparity -- as well as their interactions (Table 1). For each experiment, we compared model thresholds with those reported for human listeners with normal hearing. We quantified absolute agreement using model-to-human threshold ratios (Fig. 3a, first column) and agreement in the relative pattern of thresholds across stimulus conditions using Spearman rank correlations (Fig. 3b, first column). Across the battery, the ecologically optimized model showed close overall agreement with published group-level measures of human perceptual sensitivity. Model-to-human threshold ratios were centered near unity (median = 0.96), and model and human thresholds were strongly correlated (median ρ = 0.81, across correlations computed separately for each experiment). Results for individual experiments are examined in later sections.

**Table 1.**
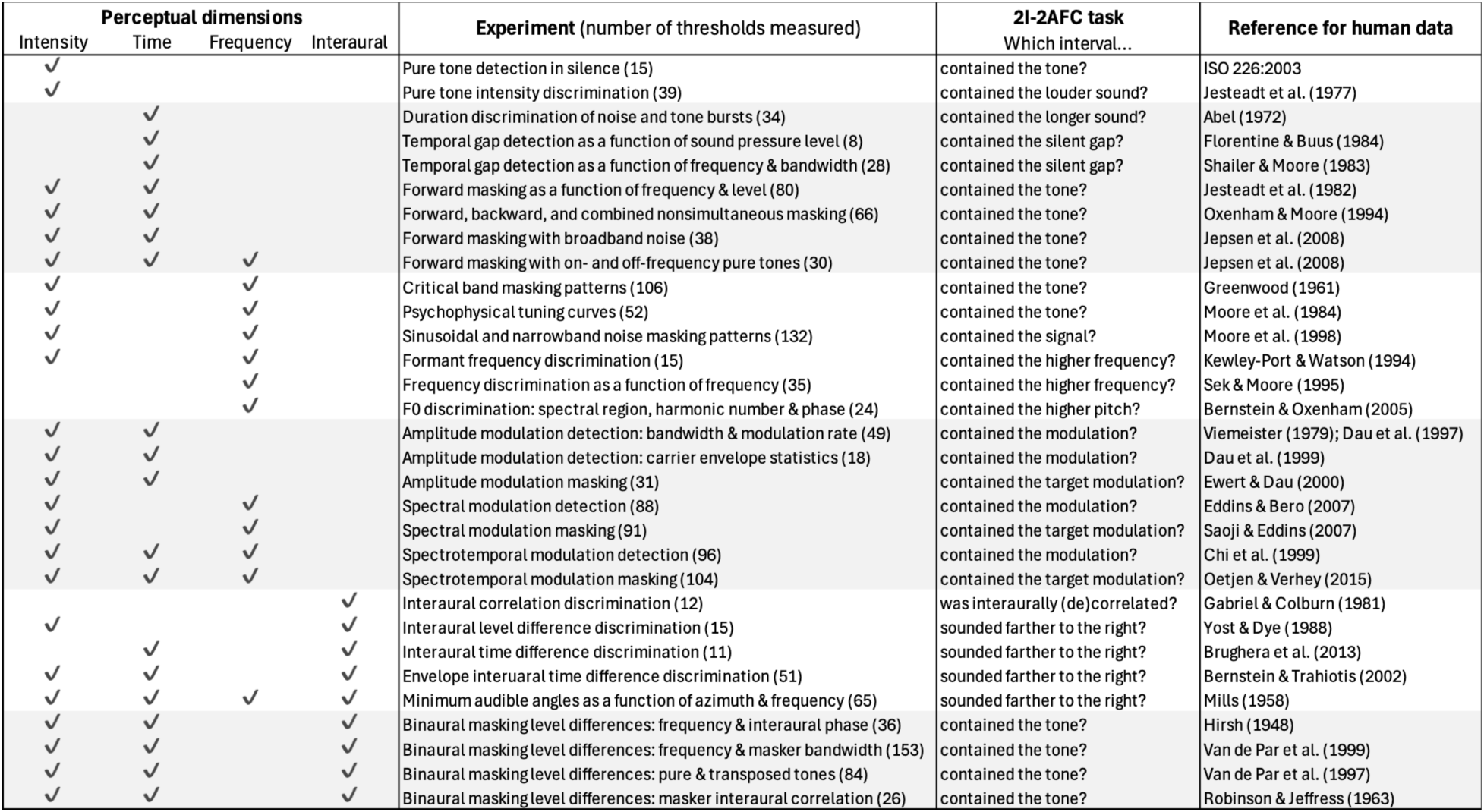
List of psychoacoustic experiments. Columns indicate the perceptual dimensions assessed, number of thresholds measured, 2I-2AFC task, and source of human data. Horizontal bands group experiments with similar dimensions or tasks: intensity, time, frequency, modulation, and binaural phenomena, as well as binaural masking level differences.

**Figure 3.**
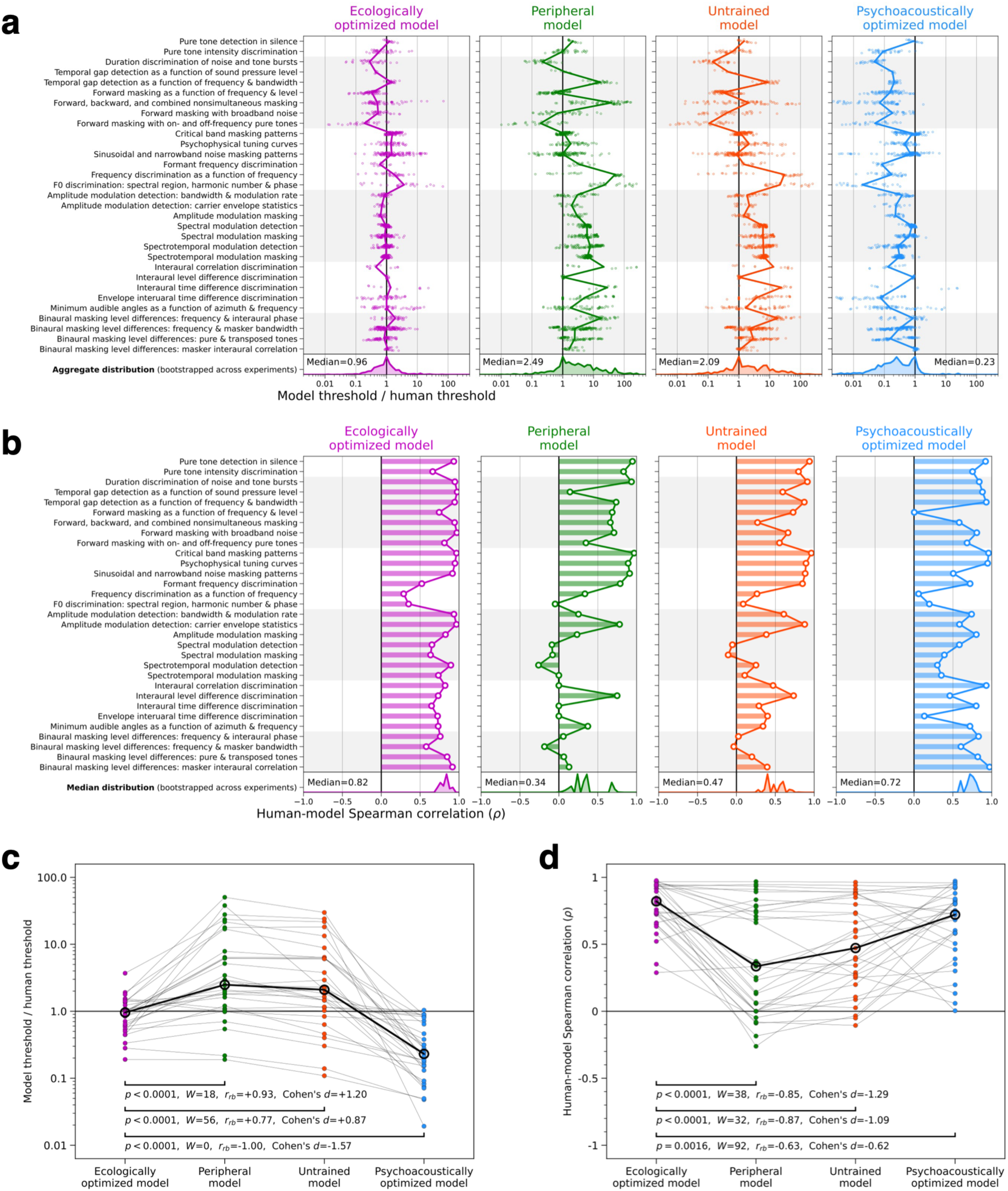
The ecologically optimized model exhibits human-like thresholds across a battery of 31 psychoacoustic experiments. (**a**) Comparison of absolute thresholds between human listeners and four model variants for each experiment. Columns show model-to-human threshold ratios computed from the model optimized for ecological hearing tasks (magenta), the model’s peripheral input representation (green), an untrained version of the model (orange), and versions of the model optimized separately for each psychoacoustic task (blue). Aggregate distributions of threshold ratios were obtained by bootstrapping threshold distributions for each experiment and combining. (**b**) Spearman correlation coefficients quantify the relative similarity between human and model thresholds across stimulus conditions within each experiment. Distributions of median correlation coefficients were obtained by bootstrapping across experiments. (**c**) Effect of model variant on median model-to-human threshold ratios. (**d**) Effect of model variant on human-model correlation coefficients. P-values and W-statistics are shown for Wilcoxon signed-rank tests comparing median values (thick lines) across the 31 experiments (thin lines). Rank biserial correlation coefficients and Cohen’s d effect sizes are shown for each comparison.

### Peripheral and untrained model representations underestimate human sensitivity

To test whether human-like thresholds required optimization for auditory tasks, we measured thresholds from two control representations: the peripheral auditory nerve representation that serves as the input to the neural network (Fig. 3, second column) and the layer 6 activations of an untrained, randomly initialized network operating on the auditory nerve input (Fig. 3, third column). The peripheral model tested the extent to which perceptual behavior could be inherited directly from the auditory periphery. The untrained model tested the contribution of the model architecture in the absence of learned representations.

Although these control models produced human-like thresholds for some simple experiments, their thresholds were substantially elevated relative to human listeners across the full battery (median model-to-human threshold ratios = 2.49 and 2.09 for the peripheral and untrained models, respectively; Fig. 3c). Their correspondence with the relative pattern of human thresholds was also substantially lower (median ρ = 0.47 and 0.34, respectively; Fig. 3d). Together, these results indicate that much of the model’s human-like psychoacoustic behavior arises from learned representations rather than from the hard-wired auditory periphery or model architecture.

### Psychoacoustically optimized model representations overestimate human sensitivity

To test whether human-like thresholds depend specifically on optimization for ecological hearing tasks, we separately optimized the network for each psychoacoustic task (Fig. 3, fourth column). Rather than holding the network weights fixed and training only a linear classifier, we jointly optimized the network and classifier weights to maximize performance on each psychoacoustic task. The weights were separately optimized for each task, creating a family of models we collectively refer to as the “psychoacoustically optimized model”. These models provide approximate lower bounds on the thresholds possible given the information available in the auditory nerve input (note that lower thresholds correspond to better performance).

The psychoacoustically optimized models substantially exceeded human performance across the battery (median model-to-human threshold ratio = 0.23). They also showed lower correspondence with the relative patterns of human thresholds (median ρ = 0.72) than the ecologically optimized model showed (Fig. 3d). These results suggest that human thresholds do not reflect optimal performance on psychophysical tasks given the information available in the sensory input. Human thresholds are better explained by the information that can be linearly decoded from representations optimized for naturalistic behavior.

### Peripheral neural stochasticity does not strongly constrain thresholds

The only source of internal noise in the model was stochastic spike sampling in the auditory nerve input representation. We included stochasticity at this stage because the auditory nerve input modeled discrete spike counts, and random sampling provided a convenient floor on the model’s absolute audibility. To assess whether this peripheral stochasticity limited perceptual sensitivity, we measured thresholds from a fully deterministic model variant. This model was optimized for the same ecological tasks using an auditory nerve representation in which sampled spike counts were replaced with the underlying expected firing rates. Eliminating spike variability makes the auditory nerve rate-level function invertible, removing the stochastic limitation on audibility. The deterministic model performed slightly better on the ecological training tasks but still closely matched human sound localization and recognition performance (Supplementary Fig. 1).

Removing peripheral stochasticity produced only modest improvements in psychoacoustic thresholds. Across the 31 experiments, the median model-to-human threshold ratio decreased from 0.96 to 0.85 (W = 91, p = 0.004; Wilcoxon signed-rank test), indicating slightly superhuman sensitivity in the deterministic model (Fig. 4a). Although statistically significant improvements were observed in 21 experiments, large effects were restricted to tasks closely tied to absolute audibility. Median thresholds improved by more than a factor of two in only two experiments: pure tone detection in silence and forward masking with long masker-signal delays. By contrast, optimizing the network representations directly for psychoacoustic tasks produced greater than two-fold improvements on 25 of the 31 experiments (Fig. 4b). Together, these results indicate that psychoacoustic sensitivity is constrained more strongly by learned representations than by peripheral stochasticity.

**Figure 4.**
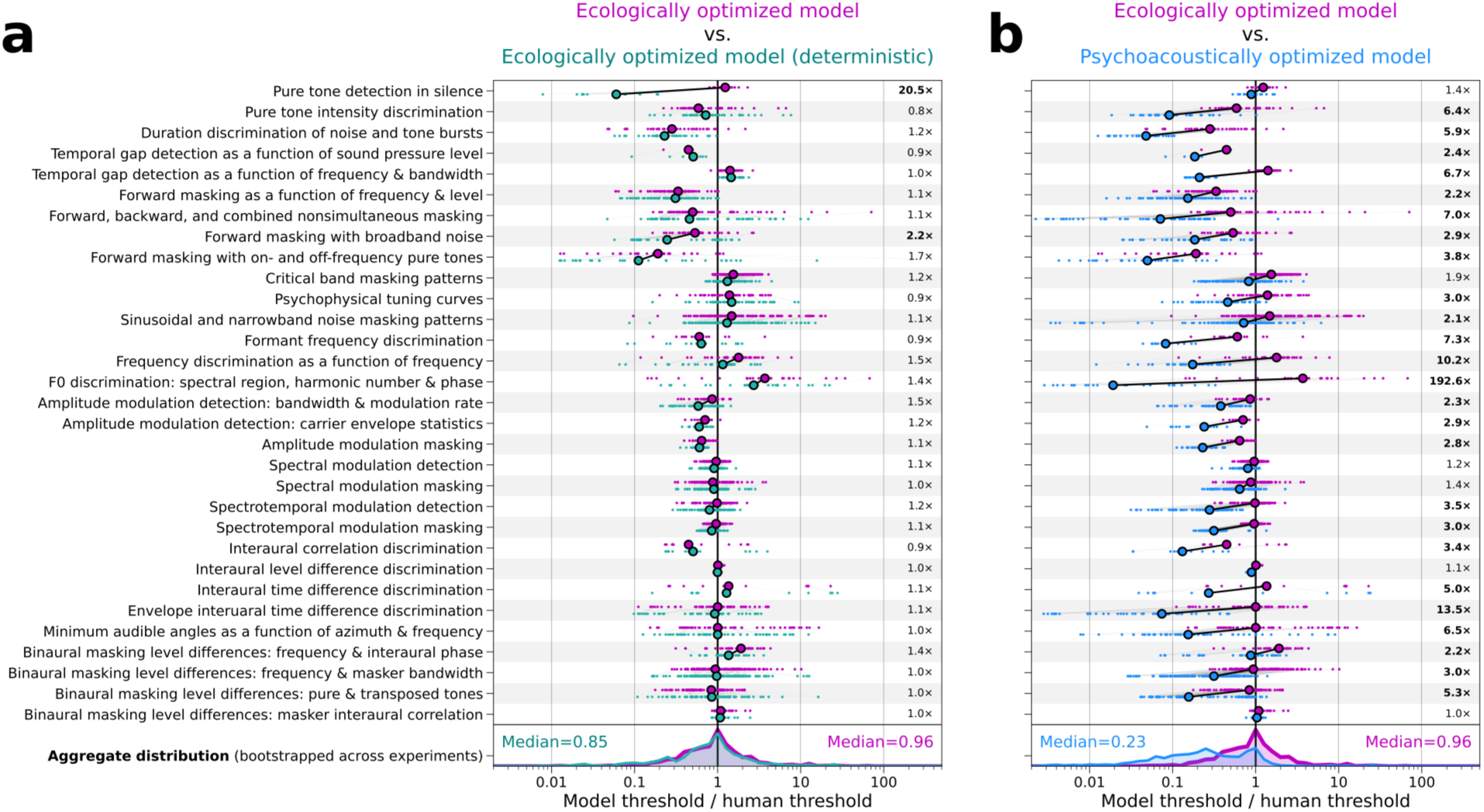
Eliminating all stochasticity from the ecologically optimized model only marginally lowers psychoacoustic thresholds. (**a**) Direct comparison of model-to-human threshold ratios between the ecologically optimized model (magenta) and a fully deterministic variant of the same model (teal). The ratio between the two models’ median thresholds is listed for each experiment, quantifying how much sensitivity improved when stochasticity was eliminated. Improvements larger than two-fold are shown in bold. (**b**) The same comparison is shown between the ecologically optimized model (magenta) and the psychoacoustically optimized model (blue). Aggregate distributions of threshold ratios were obtained by bootstrapping threshold distributions for each experiment and combining.

### Human-like modulation sensitivity emerges from optimization for ecological hearing tasks

Having characterized model behavior across experiments in aggregate, we next examine several representative paradigms in detail. Because natural sounds contain fluctuations over time, frequency, and both dimensions jointly, human sensitivity to these fluctuations, quantified as thresholds for detecting or discriminating modulations along either or both dimensions (Fig. 5a), has long provided an important benchmark for theories of auditory processing^4,29,30^.

**Figure 5.**
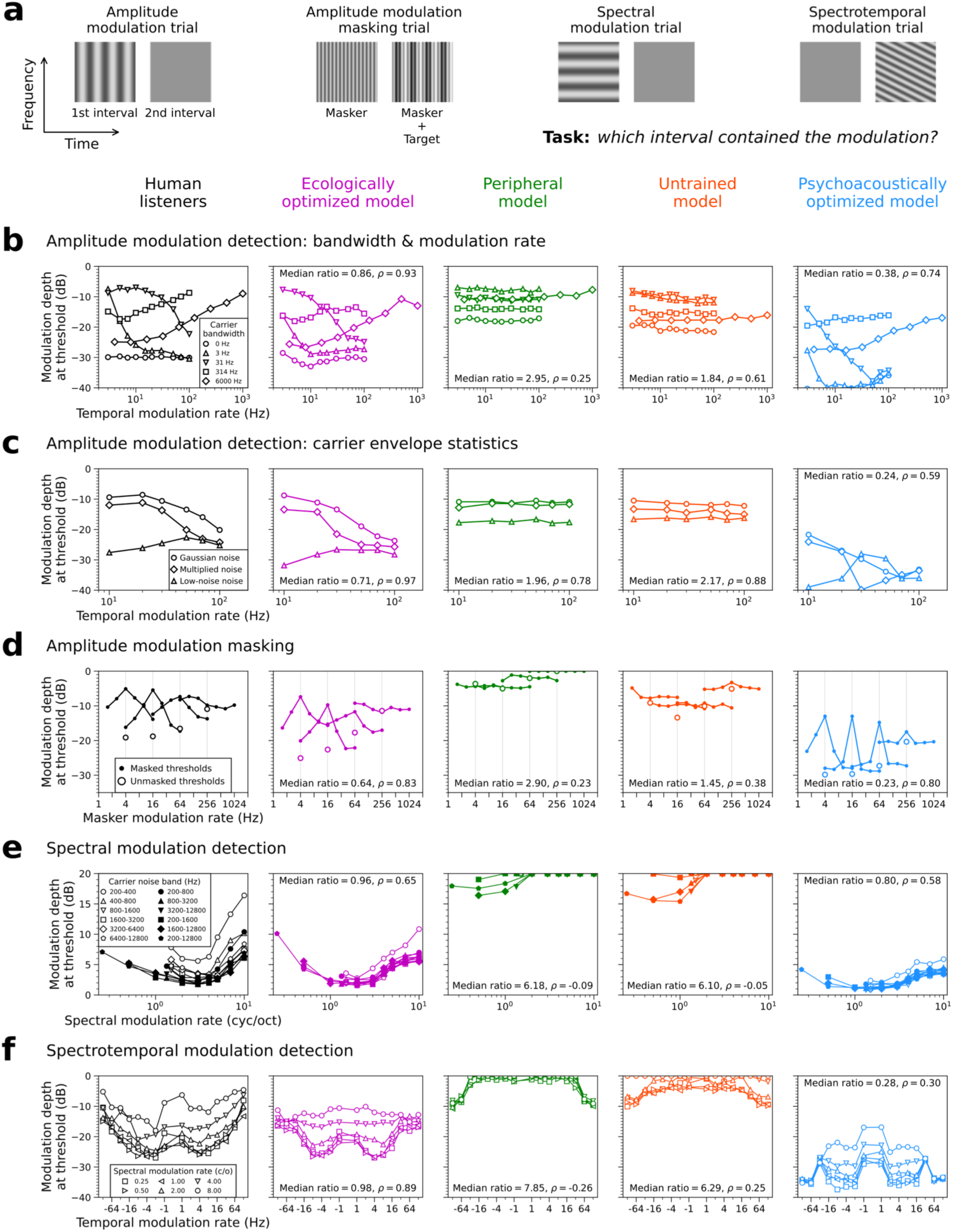
Ecologically optimized model representations support human-like modulation sensitivity. (**a**) Schematic spectrograms illustrating trials for amplitude modulation detection, amplitude modulation masking, spectral modulation detection, and spectrotemporal modulation detection tasks. Panels (**b**-**f**) are organized into five columns, showing thresholds measured from human listeners and four model variants: the ecologically optimized model, peripheral model, untrained model, and psychoacoustically optimized model. (**b**) Amplitude modulation detection thresholds measured as a function of temporal modulation rate and carrier bandwidth. (**c**) Amplitude modulation detection thresholds measured for three types of carrier noise with the same bandwidth but different temporal envelope spectra. (**d**) Amplitude modulation detection thresholds measured in the presence of different masker modulations. The modulation rate of the target was 4, 16, 64, or 256 Hz. The modulation rate of the masker varied across trials. (**e**) Spectral modulation detection thresholds measured as a function of spectral modulation rate and carrier noise band. (**f**) Spectrotemporal modulation detection thresholds measured as a function of temporal and spectral modulation rate. The sign of the temporal modulation rate determines whether the spectrotemporal modulation is oriented up or down. The spectrotemporal modulation depicted in (**a**) has a negative temporal modulation rate. Median model-to-human threshold ratios and human-model Spearman correlation coefficients (ρ) are computed across the stimulus conditions within each experiment.

We first measured amplitude modulation detection thresholds by training linear classifiers to discriminate model representations of sinusoidally amplitude-modulated and unmodulated noise in a 2I-2AFC task. Thresholds were measured as a function of modulation rate for noise carriers with different bandwidths^4,29^ (Fig. 5b) and for noise carriers with the same bandwidth but different temporal envelope spectra^31^ (Fig. 5c). In each case, thresholds were measured from four model variants presented in the previous sections: the ecologically optimized model, the peripheral model, the untrained model, and the psychoacoustically optimized model. Despite never being fit to human data, the ecologically optimized model produced close-to-human thresholds across modulation rates and the different noise carriers.

We next examined amplitude modulation masking, measuring the detectability of a target modulation in the presence of masker modulations that varied across trials^32^. In humans, this paradigm reveals tuning to modulation frequency, with thresholds highest when the masker and target modulation are similar (Fig. 5d). The ecologically optimized model produced similarly tuned masking functions that resembled human measurements.

The correspondence extended to spectral and spectrotemporal modulation. Spectral modulation detection thresholds vary systematically with spectral modulation rate and carrier noise band in humans^33^, and these trends were reproduced by the ecologically optimized model (Fig. 5e). Likewise, the ecologically optimized model reproduced patterns of human sensitivity to spectrotemporal modulations across different combinations of spectral and modulation rate^34^ (Fig. 5f). The ecologically optimized model also reproduced human patterns of spectral^35^ and spectrotemporal^36^ modulation masking (Supplementary Fig. 2).

The agreement with human behavior in these experiments was specific to the ecologically optimized model. Untrained and peripheral model representations underestimated human sensitivity and failed to reproduce characteristic patterns of modulation selectivity. By contrast, representations optimized directly for modulation detection overestimated human sensitivity and exhibited superhuman modulation selectivity. Together, these results suggest that the characteristic limits of human modulation sensitivity emerge from perceptual representations optimized for everyday hearing.

### Human-like binaural sensitivity emerges from optimization for ecological hearing tasks

Human spatial hearing depends on sensitivity to interaural level differences (ILDs) and interaural time differences (ITDs) as well as computations that integrate these cues across frequency and time. We examined a series of psychoacoustic paradigms probing both elementary binaural sensitivity and more complex binaural interactions (Fig. 6).

**Figure 6.**
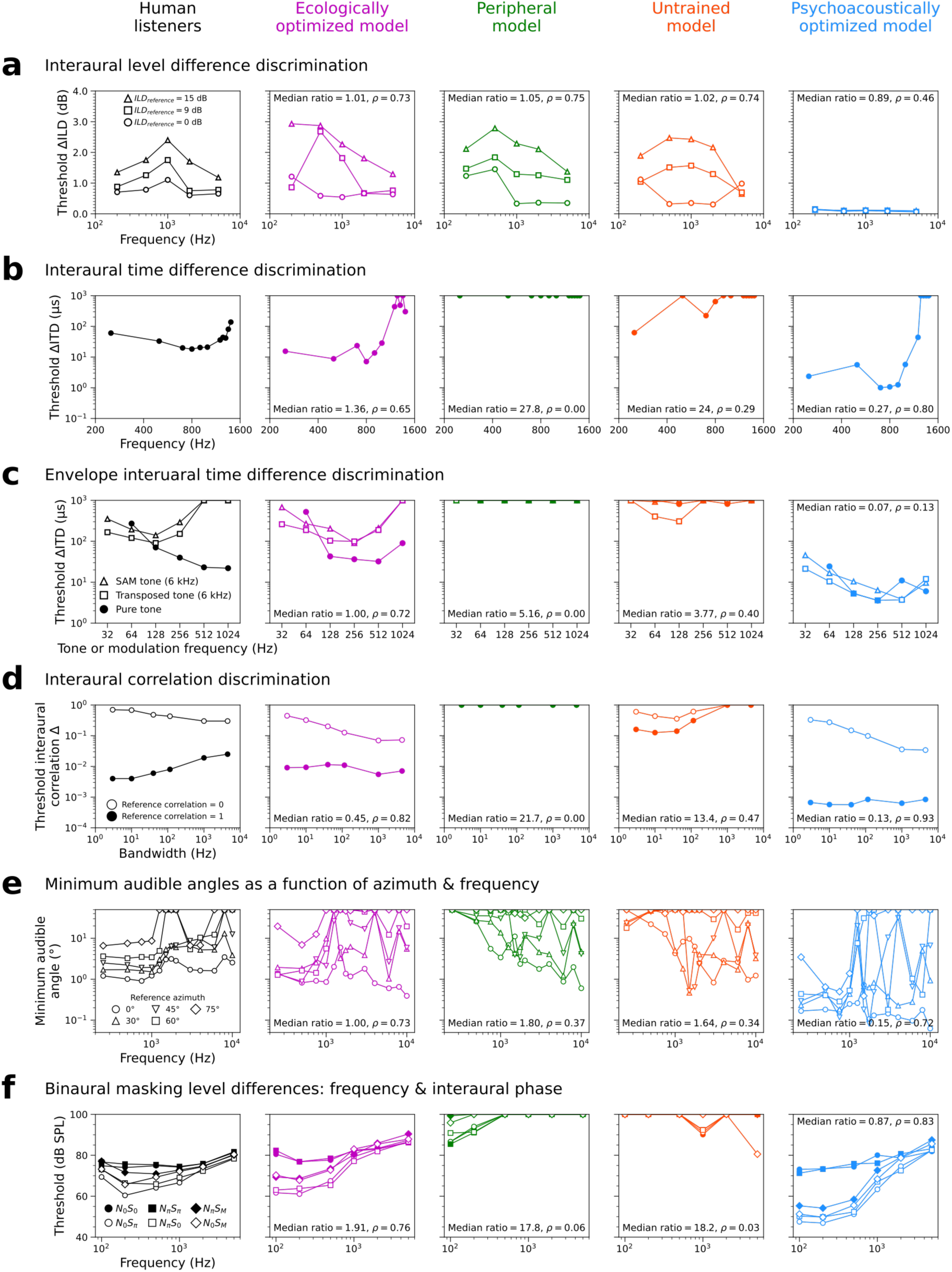
Ecologically optimized model representations support human-like binaural cue sensitivity. Results are organized into five columns plotting thresholds measured from human listeners and four model variants: the ecologically optimized model, the peripheral model, the untrained model, and the psychoacoustically optimized model. (**a**) Pure tone ILD discrimination thresholds measured as a function of frequency at three reference ILDs. (**b**) Pure tone ITD discrimination thresholds measured as a function of frequency. (**c**) Envelope ITD discrimination thresholds measured as a function of modulation frequency for sinusoidally amplitude-modulated (SAM) tone and transposed tones. ITD thresholds for pure tones are included on the same axes for comparison. (**d**) Interaural correlation discrimination thresholds measured as a function of noise bandwidth at two reference interaural correlations. When the reference correlation is 0, the task is to detect which interval is more correlated between the two ears. When the reference is 1, the task is to detect which interval is less correlated. (**e**) Minimum audible angles measured as a function of azimuth and frequency. (**f**) Binaural tone-in-noise detection thresholds measured as a function of tone frequency and interaural phase condition. The interaural phase of the noise (N) and signal (S) are each denoted with subscripts: 0 = in phase between the ears, π = 180° out of phase, and M = monaural presentation. Median model-to-human threshold ratios and human-model Spearman correlation coefficients (ρ) are computed across the stimulus conditions within each experiment.

The ecologically optimized model reproduced the qualitative dependence of ILD (Fig. 6a) and ITD (Fig. 6b) discrimination thresholds on stimulus frequency, with absolute thresholds generally comparable to those of human listeners^37,38^. The model also exhibited qualitatively human-like envelope ITD sensitivity (Fig. 6c), although thresholds at high modulation frequencies were considerably lower than those measured in humans^39^. The model’s interaural correlation discrimination thresholds across noise bandwidths (Fig. 6d) and minimum audible angle thresholds as a function of stimulus frequency and azimuth (Fig. 6e) were also generally consistent with those of human listeners^40,41^. Across these experiments, peripheral and untrained representations substantially underestimated human sensitivity, whereas psychoacoustically optimized representations exhibited superhuman performance. The one exception was the ILD experiment (Fig. 6a), where the ecologically optimized, untrained, and peripheral model representations produced similarly human-like thresholds.

Binaural masking level differences (BMLDs) are a well-documented phenomenon in which changing the interaural phase relationship of a tone signal or noise masker substantially alters tone detection thresholds in noise^42^. Because BMLDs have long served as a benchmark for mechanistic theories of binaural hearing^3^, they provide a useful test of the present framework. The ecologically optimized model reproduced the principal patterns of human BMLDs across signal frequencies^42^ (Fig. 6f), masker bandwidths^43^, carrier types^44^, and masker interaural correlations^45^ (Supplementary Fig. 3). By contrast, peripheral and untrained representations failed to produce appreciable BMLDs, whereas psychoacoustically optimized representations consistently overestimated their magnitude. Altogether, these results indicate that representations optimized for everyday hearing tasks capture many characteristic aspects of human binaural perception, although some quantitative differences remain for specific measures of binaural sensitivity.

### Ecologically optimized model exhibits human-like behavior on many but not all psychoacoustic experiments

Across the remaining psychoacoustic experiments, the ecologically optimized model also produced a strikingly human-like pattern of behavior (Fig. 7). Thresholds generally tracked human measurements across a broad range of frequency, intensity, and temporal conditions, encompassing intensity discrimination^46^, duration discrimination^47^, gap detection^48,49^, nonsimultaneous masking^50–52^, frequency selectivity^53–55^, frequency discrimination^56,57^, and pitch discrimination^58^ paradigms. Here, we highlight the notable deviations from human behavior.

**Figure 7.**
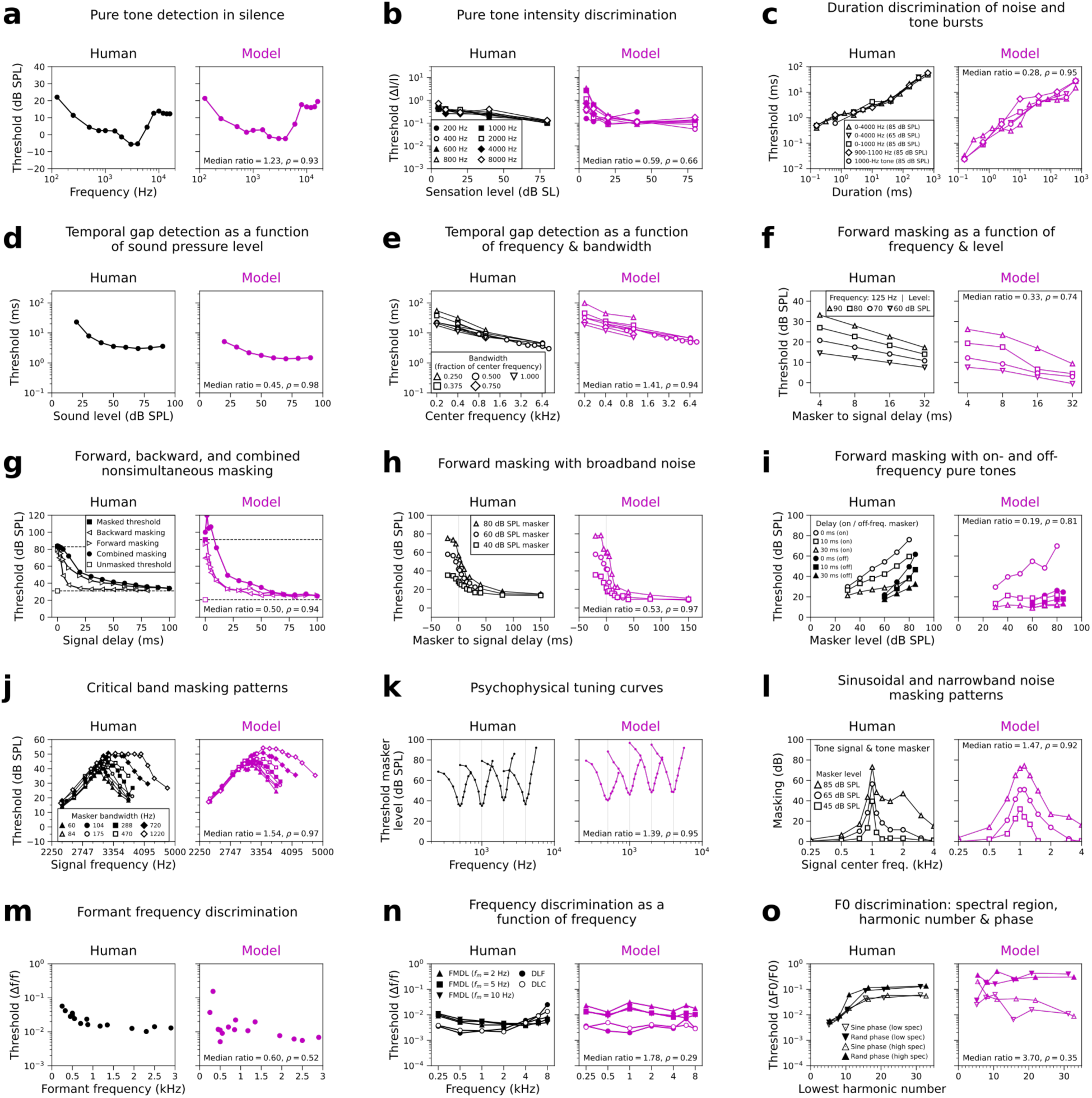
Ecologically optimized model representations produce human-like behavior in most of the remaining experiments in the battery. For each experiment (**a-o**), thresholds from human listeners and the ecologically optimized model are shown as a function of the same stimulus parameters. Median model-to-human threshold ratios and human-model Spearman correlation coefficients (ρ) are computed across the stimulus conditions within each experiment. Notable discrepancies between human and model behavior occur in six panels. In (**c**), the model exhibits superhuman duration discrimination for stimuli less than 10 ms in duration. In (**g**), the model does not reproduce the asymmetry between forward and backward masking. In (**i**), the model poorly matches the different slopes produced by on- and off-frequency forward maskers. In (**l**), the model fails to capture level-dependent masking patterns. In (**n**), the model’s difference limens for frequency discrimination (DLF) and frequency change detection (DLC) do not exhibit a human-like a dependence on frequency. In (**o**), the model’s F0 discrimination thresholds do not exhibit a human-like dependence on lowest harmonic number.

The model exhibited superhuman duration discrimination for short-duration stimuli (Fig. 7c). In nonsimultaneous masking experiments, the model reproduced the general dependence of thresholds on temporal separation between signal and masker but did not capture several qualitative characteristics of human performance. In particular, the model failed to reproduce the asymmetry between forward and backward masking (Fig. 7g) and underestimated the magnitude of forward masking for high-frequency pure tones (Supplementary Fig. 4). These discrepancies may reflect limitations of the model’s treatment of time: the network uses convolutional filters operating over time and frequency rather than dynamics that unfold causally over time. The model also failed to reproduce the qualitative effects of on- and off-frequency forward maskers (Fig. 7i) and level-dependent simultaneous masking patterns (Fig. 7l and Supplementary Fig. 5). These effects are typically attributed to frequency-dependent, non-linear cochlear compression^59^, which was absent from the simplified peripheral model used here.

Fine frequency and pitch judgments revealed additional deviations. Pure tone frequency discrimination thresholds were generally comparable to those of human listeners, but the model did not reproduce the characteristic increase in thresholds at frequencies above ∼3 kHz (Fig. 7n), a pattern often interpreted as reflecting a transition from temporal fine structure to place-based coding^57,60^. Similarly, F0 discrimination thresholds for harmonic complex tones were comparable to human thresholds for stimuli containing only high-numbered unresolved harmonics, but the model did not exhibit the marked improvement for stimuli containing resolved harmonics observed in humans^61^ (Fig. 7o). This discrepancy is consistent with the possibility that the model does not exploit temporal fine structure cues^20^, presumably because the set of training tasks we used do not require it to do so.

For many experiments in Figure 7, similar performance was obtained using the peripheral representation alone (Supplementary Figs. 4-6), indicating that many of the behaviors were largely inherited from the auditory nerve input rather than emerging from learned features. By contrast, psychoacoustically optimized representations exhibited superhuman sensitivity in nearly all the experiments, indicating that greater sensitivity is possible given the peripheral input, but it is not achieved by the ecologically optimized model.

### Emergence of human-like psychoacoustic behavior across the network hierarchy

Although previous work established a coarse correspondence between the layers of similar neural network models and putative stages of the auditory cortex^16,62^, it is nevertheless informative to ask where human-like psychoacoustic behavior emerges in the model’s hierarchy of processing stages. The thresholds analyzed in Figures 3-7 were measured from the deepest convolutional block (layer 6) shared across the ecological training tasks. To evaluate this choice, we independently trained linear classifiers for each psychoacoustic task using activations from each network stage. For stages beyond the network’s branch point, activations from the four task-specific branches were concatenated.

We compared thresholds measured from each model stage to those of humans across the full battery of experiments. Human-model agreement generally improved deeper into the model, with median model-to-human threshold ratios converging to unity (Fig. 8a) and median human-model rank correlations increasing across stages (Fig. 8b). Most of these improvements occurred before layer 4, after which performance remained comparatively stable despite substantial variation across individual psychoacoustic tasks. Layer 6 falls within this stable regime of human-like performance, supporting its use as the default representation throughout this work.

**Figure 8.**
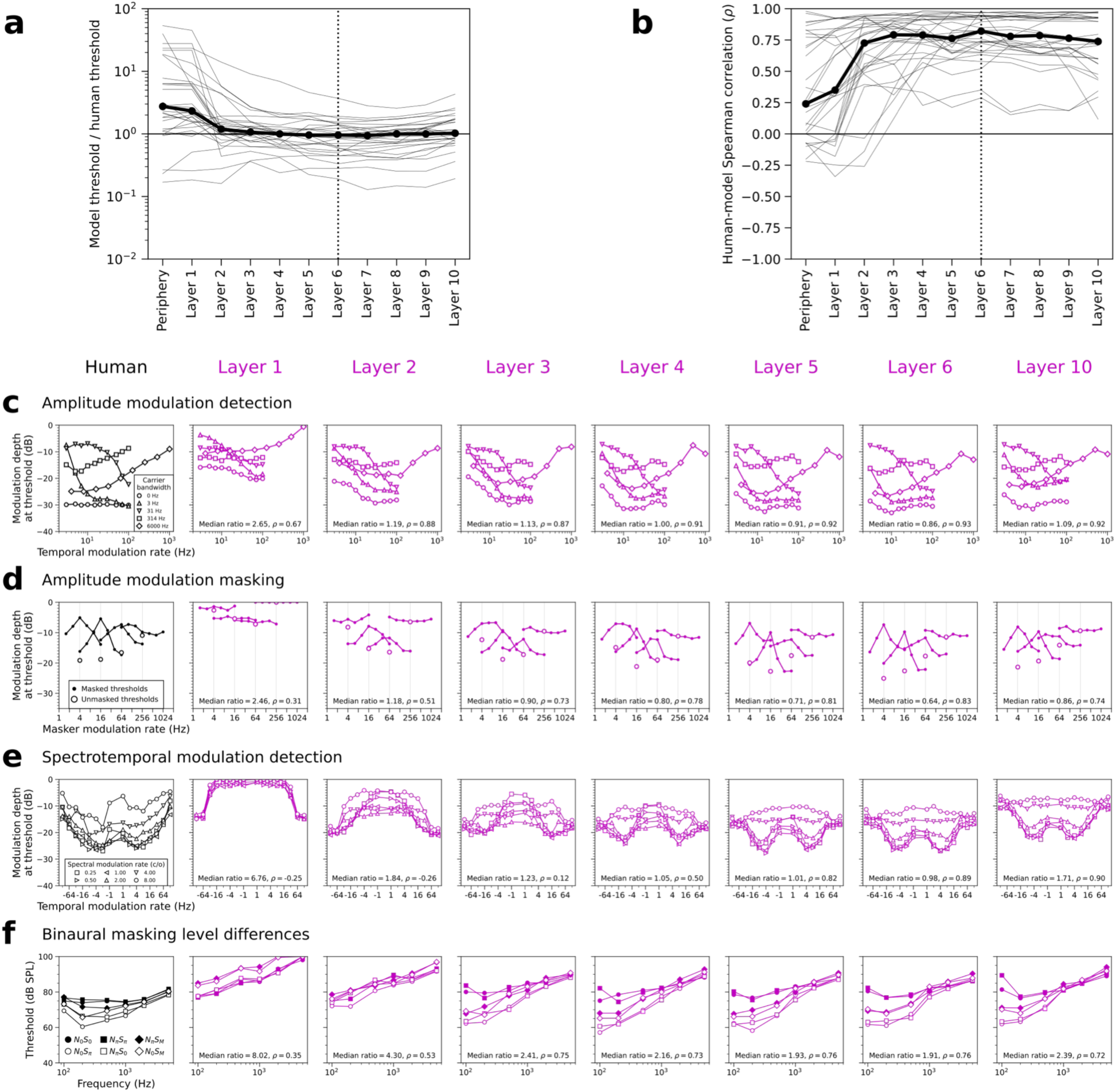
Emergence of human-like psychoacoustic sensitivity across the network hierarchy. The ecologically optimized model’s psychoacoustic thresholds were separately measured from each model stage. (**a**) Model-to-human threshold ratios were computed for each of the 31 experiments (thin lines) as a function of model layer. The thick line indicates the median across experiments. The dashed vertical line denotes the default layer used in previous figures. (**b**) Human-model Spearman correlation coefficients for each experiment as a function of model layer. (**c**) Amplitude modulation detection thresholds measured from human listeners (in black) and seven model stages (in magenta). Thresholds are similarly shown for the amplitude modulation masking (**d**), spectrotemporal modulation detection (**e**), and binaural masking level difference (**f**) experiments. Median model-to-human threshold ratios and human-model Spearman correlation coefficients (ρ) are computed across the stimulus conditions within each experiment.

Specific psychoacoustic traits emerged at different stages of the network hierarchy. Sensitivity to temporal and spectrotemporal modulations (Fig. 8c-e) became increasingly human-like through layer 5, suggesting this behavior emerges progressively across multiple stages of hierarchical processing. Most binaural phenomena first appeared at layer 2, the earliest stage at which information from the two ears was combined (the first convolutional stage processes the two ears independently to loosely mirror the organization of the mammalian auditory pathway). Binaural masking level differences became larger and more human-like following binaural integration (up to layer 4; Fig. 8f), suggesting these phenomena also emerge progressively across the network hierarchy.

## DISCUSSION

We developed an auditory model optimized for naturalistic listening behavior to test whether such optimization could account for human perceptual sensitivity. The model jointly learned to localize and recognize words, voices, and environmental sounds from simulated auditory nerve representations of natural scenes. We then measured a broad range of perceptual thresholds using classifiers operating on the model’s learned representations. The model exhibited human-like psychophysical thresholds across many different stimulus dimensions, demonstrating that absolute thresholds can be largely explained by the linear separability of stimuli within representations optimized for ecological tasks. Thus, human perceptual sensitivity is as good, and only as good, as it needs to be in the service of natural behavior.

A common explanation for absolute psychophysical thresholds is that they are constrained by internal noise arising from the stochastic nature of biological hardware^3,5,6^. Here, however, the model produced human-like thresholds without requiring sources of internal noise beyond stochasticity in the auditory nerve representation. Removing even this peripheral stochasticity produced only modest improvements in thresholds across most experiments. Conversely, optimizing the same model directly for individual psychoacoustic tasks produced substantially superhuman sensitivity, demonstrating that the model’s human-like thresholds were not simply a consequence of limited representational capacity. Together, these results suggest that internal noise is not needed to explain human perceptual limits, and that representations optimized for ecological behavior are sufficient to account for most aspects of auditory sensitivity. This conclusion is consistent with proposals that deep learning instantiates efficient codes^24^, and with theories that efficient codes constrain discriminability^14,15^, but provide a much more comprehensive test of such ideas than was previously possible.

Traditional models of psychoacoustic phenomena typically begin with hypothesized mechanisms designed to explain individual effects. Such models have provided important insights into auditory processing, but they generally do not address why these mechanisms should exist or whether they are sufficient to support natural listening behavior. Here, many classic psychoacoustic phenomena emerged in a model trained only to perform natural listening tasks. Previous work has shown that artificial systems optimized for sensory tasks reproduce aspects of human perceptual sensitivity^17,19,20,24,63,64^. The present work extends this evidence by demonstrating such correspondence across a comprehensive set of psychophysical phenomena. Our results support a normative account of auditory perception in which characteristic patterns of human sensitivity arise because the auditory system is optimized for the demands of everyday behavior.

### Human-model discrepancies

The model’s failures hint at constraints that plausibly shape auditory perception but that were missing from the present model’s optimization. The largest discrepancies involved fine frequency and pitch judgments, for which the model did not reproduce qualitative patterns commonly associated with temporal fine structure processing^57,60^. One possibility is that the ecological training tasks used here did not sufficiently capture the demands shaping human pitch perception. For example, the model was not optimized for tasks involving music or multi-talker situations, both of which may place stronger demands on pitch processing. Alternatively, these discrepancies may reflect limitations of the network architecture, which may not efficiently extract temporal fine structure from the auditory nerve representation.

Although the model reproduced many aspects of human binaural hearing, it overestimated sensitivity to high-frequency envelope ITDs and did not fully capture all patterns of binaural masking level differences. These discrepancies may reflect biological constraints on binaural computations that were neglected in the present model. For example, the optimization objective did not account for the wiring or metabolic costs^65^ associated with specialized binaural circuitry, which could plausibly shape binaural representations.

As noted above, discrepancies in temporal sensitivity, likely reflect limitations of the feed-forward convolutional architecture, which lacks explicit internal dynamics unfolding over time. Likewise, failures to reproduce some level- and frequency-dependent masking effects are consistent with the simplified linear cochlear filtering used in the model’s periphery^52,59^. Thus, some of the model’s deviations from human behavior point to identifiable biological or architectural constraints that could be incorporated into future models.

### Limitations

Our results do not establish that ecological task optimization is uniquely responsible for human-like thresholds. Other forms of optimization over natural sounds, including self-supervised objectives, or potentially objectives that explicitly incorporate measures of efficiency, might produce similarly human-like perceptual sensitivity. Determining which aspects of the optimization objective, natural stimulus statistics, and architectural constraints are critical for producing human-like sensitivity remains an important question for future work.

Although the psychoacoustic battery we compiled spanned the principal dimensions of auditory nerve representations, it does not encompass all aspects of hearing or every possible interaction between dimensions. For instance, interactions between frequency and interaural disparity were relatively underrepresented, in part because many binaural cue comparisons are thought to occur within frequency channels^66^.

Human thresholds were compiled from the psychoacoustic literature, and as a result represent the average results for a group of participants. These measurements do not capture the individual differences that are typically present in groups of human listeners due to differences in experience^67^, hearing health^68^, and other factors. Future work using larger datasets obtained from the same listeners across many tasks, along with models that capture some of the plausible sources of between-listener variance, could provide a way to shed light on individual differences.

We assumed linear decision rules. Such rules provide a straightforward and standard measure of the explicitly available information in learned representations^69–72^ but could underperform nonlinear decision rules. If humans rely on nonlinear decision rules, internal noise might be needed to explain perceptual sensitivity (by offsetting the benefits of a more expressive decoder). Our results show that it is possible to account for human thresholds without appealing to internal noise, but we cannot rule out the possibility that human thresholds reflect a combination of nonlinear decision rules and internal neural noise.

It also remains possible that the results are somewhat specific to the size of the model architecture and the scale of the training data it was optimized for. We chose an architecture and training data that are sufficient to account for human performance of ecological tasks and found that the resulting model also explains perceptual sensitivity. However, larger models trained on vast amounts of data could potentially exhibit superhuman performance on ecological tasks, and potentially also better perceptual sensitivity.

As in other large neural network models, the mechanisms underlying the model’s behavior are not readily interpretable. The layer-wise analyses provided some insight into where different psychoacoustic phenomena emerge across the network hierarchy, but many behaviors developed progressively over multiple stages rather than from easily identifiable computational components. Developing a more mechanistic understanding may require incorporating model stages that explicitly implement hypothesized computations.

### Future directions

Models like those used here but with simulated hearing loss in the auditory nerve representation could clarify how peripheral impairments translate to altered perceptual limits^73–76^. Such predictions may help relate audiological measurements to real-world hearing abilities and guide the development of targeted compensation strategies.

Our results raise the question of how stochasticity in the brain relates to behavior^77,78^. There is considerable evidence for some irreducible sources of noise in the nervous system^7^. At the same time, many components of trial-to-trial variability in neural responses are argued to represent effects of state rather than noise per se^79,80^, in which case they might not be expected to limit behavior^81^. The ability to build models of real-world behaviors offers a new avenue to understand the role of stochasticity in neural computation and behavioral abilities.

The approach developed here provides a general method for relating artificial sensory systems to human psychophysical abilities. By training linear classifiers to make psychophysical judgments using model representations, model thresholds can be directly compared with human thresholds. Although applied here to the sense of hearing, this approach is equally applicable to other sensory domains.

## METHODS

### Training dataset for ecological hearing tasks

We trained the model on 7,644,413 unique 2.2-s binaural auditory scenes (sampled at 44.1 kHz) designed to capture aspects of everyday hearing. Scenes contained a speech or an environmental-sound foreground presented at a single spatial location, embedded in spatially distributed non-speech background sounds. Speech foregrounds were drawn from Common Voice^82^ and nonspeech foregrounds were drawn from the GISE-51^83^ subset of FSD50K^84^. Backgrounds were drawn from Audio Set^85^. 80% of scenes contained speech foregrounds and 20% contained environmental sound foregrounds. Foregrounds were spatialized in 90% of scenes and presented diotically in the remainder, providing exposure to both spatialized and headphone-presented stimuli. Backgrounds were rendered from 1 to 9 spatial locations for speech scenes and 3 to 9 locations for environmental-sound scenes. In 5% of scenes, the foreground was presented without background noise; otherwise, SNRs were uniformly sampled from -10 to +15 dB, with individual background sources independently level-roved by ±2.5 dB before summation. Foreground levels were sampled uniformly between 30 and 90 dB SPL.

The dataset was spatialized using KEMAR HRTFs^86^ simulated in 2000 acoustically distinct shoebox rooms (1800 for training, 200 for validation)^17^. Rooms varied in size and shape, acoustic absorption, and listener position and orientation. For each room, binaural room impulse responses were simulated via the image-source method^87^ for 72 azimuths (0 to 360° in 5° steps), 7 elevations (0 to 60° in 10° steps), and two source distances: one fixed at 1.4 m and one randomly sampled between 1 m and 31 m. Foreground and background clips were sampled with replacement as needed to balance class labels while maximizing the number of unique sources in the final dataset. To increase variability of the source clips, randomly sampled band-pass filters (2 to 4 octaves wide) were applied to 50% of the foregrounds. Small shifts in F0 (up to ±0.5 semitones), tempo (up to ±20%), and speed (up to ±2%) were randomly sampled and applied to 25%, 12.5%, and 12.5% of the speech foregrounds, respectively, and using SoX audio transformations.

Scenes were labeled for four classification tasks: foreground location (504 azimuth-elevation pairs), middle-word recognition (809 words), voice recognition (500 familiar talkers), and environmental sound classification (49 classes). Each task also included a null label for stimuli for which the corresponding task was undefined or outside the model’s target distribution. The 809-word vocabulary was selected to maximize the number of unique female talkers represented and included the digits 0 to 9. The 500 familiar talkers (250 female and 250 male) produced 43% of the speech utterances in the final dataset, which included clips from 51,870 unique talkers (38% female, 61% male, and 1% nonbinary or not reported). The 49 environmental sound classes were the category labels from GISE-51, excluding “Human speech” and “Whispering”.

### Auditory nerve input representation

We used the simplified auditory nerve model of Saddler & McDermott (2024), which has previously produced deep neural network models whose behavior closely matches that of models operating on more detailed auditory nerve representations^17,88^. Audio was resampled to 50 kHz and passed through a static middle-ear/audibility filter matched to ISO 226:2003 thresholds, followed by a 50-channel ERB-spaced gammatone filterbank spanning 60 to 16,000 Hz, implemented using a 50-ms finite impulse response approximation. Filter outputs were half-wave rectified, low-pass filtered at 3 kHz, and downsampled to 10 kHz. Auditory nerve firing rates were then computed using sigmoid rate-level functions defined on a dB scale for high-, medium-, and low-spontaneous-rate (HSR, MSR and LSR) fibers. HSR, MSR, and LSR rate-level functions had thresholds of 0, 12, and 28 dB SPL, dynamic ranges of 20, 40, and 80 dB, and firing rate ranges of 70 to 250, 4 to 250, and 0.1 to 100 spikes per second, respectively. We represented the spiking activity of 32,000 auditory nerve fibers per ear, comprising 60% HSR, 25% MSR, and 15% LSR fibers^89^. At each time point, the summed spike count for each fiber type and frequency channel was sampled independently using the normal approximation to the binomial distribution. Refractory effects were neglected because the representation pooled the activity of 96 to 384 fibers within each frequency channel. For stimuli longer than 2 s, a random 2-s excerpt was selected after generating the full auditory nerve response. The resulting input representation was a 6 x 50 x 20,000 tensor of auditory nerve spike counts, with the six channels corresponding to the three fiber types in each ear. Spike count sampling was the sole source of internal noise in the model. We additionally evaluated fully deterministic models in which sampled spike counts were replaced by their binomial expectations, computed as the number of fibers multiplied by the instantaneous spiking probabilities.

### Neural network architecture

Auditory nerve representations were processed by a branched feed-forward convolutional neural network containing 685,134,090 learnable parameters. Six convolutional blocks formed a shared trunk, after which processing diverged into four task-specific branches. Each block comprised two-dimensional convolution, SiLU activation, Hanning-window-weighted average pooling, and layer normalization. In the first block, the convolution was divided into two groups to process the left- and right-ear auditory nerve representations separately^17^, loosely mirroring the initial separation of binaural inputs in the auditory system. Subsequent blocks used standard convolutions to integrate information across ears. The task-specific branches contained four additional convolutional blocks followed by a 512-unit fully connected layer, SiLU activation, layer normalization and 50% dropout during training. The architecture was obtained by iterative refinement of an architecture previously identified in a large-scale random architecture search^90^. The branch point was placed as deep as possible into the network without impairing individual task performance^16^, although this criterion was formally evaluated using an earlier version of the architecture. Although here we present results from only one network neural network architecture, we note that pilot experiments with earlier iterations of the model with different architectures and initializations produced very similar behavior and perceptual thresholds.

### Neural network optimization

The network was trained to minimize the summed softmax cross-entropy losses across the four tasks, with task weights sampled independently at random for each example and null-labeled tasks excluded from the loss. Parameters were optimized using Adam with a learning rate of 1e-5 and a batch size of 128 across four NVIDIA H100 GPUs. Training continued for 500,000 steps (16 million examples; approximately 70 h) until performance had converged across tasks, as assessed from validation curves. Model checkpoints were evaluated every 2,500 steps. For the localization, word recognition, and voice recognition tasks, validation accuracies were rank-ordered and the checkpoint with the lowest summed rank was selected. The environmental sound recognition task was excluded from checkpoint selection because its validation accuracy plateaued substantially earlier.

### Model evaluation: ecological task performance

After training, all network weights were frozen, and we refer to this trained model as the “ecologically optimized model”. We compared its performance with normal hearing human listeners on five naturalistic tasks: sound localization in noise, word recognition in real-world noise, word and voice recognition with F0-shifted speech, environmental sound recognition, and spatial release from masking. Human data were taken from previous experiments, and the corresponding model stimuli and task procedures were matched to the human experiments as closely as possible.

#### Natural sound localization in noise

We compared the model with human sound localization in noise data from ref.^17^, in which 11 listeners (21 to 30 years; 5 female, 6 male) localized 460 unique GISE-51 environmental sounds presented from a 19 x 5 hemispherical loudspeaker array (2-m radius; 19 azimuths spanning 0 to 180° and 5 elevations spanning 0 to 40°, in 10° increments). Each 1-s target (44.1 kHz; including 15-ms half-Hanning onset and offset ramps) was presented at 60 dB SPL (A-weighted) from one of the 95 loudspeakers while threshold-equalizing noise played concurrently from nine other loudspeakers, chosen at random. Noise levels were scaled to produce target SNRs of -13.6, -6.8, 0, +6.8 or +13.6 dB on each trial. Listeners reported the target loudspeaker without moving their heads during sound presentation. We presented the same sounds in a virtual rendering of the loudspeaker array room and evaluated the model’s localization judgments across all 460 sounds, 95 loudspeaker locations, and 5 SNRs (218,500 stimuli). For the model, 1-s targets were centered in continuous 2-s background noise to ensure the localization task was well-defined. Model localization judgments were limited to the 95 speaker locations to match task difficulty. We compared mean absolute spherical (great circle), azimuth, and elevation errors between model and human responses as a function of SNR.

#### Word recognition in noise

We compared the model with human word-in-noise recognition data from ref.^17^, collected from self-reported normal hearing listeners across two online experiments. In both experiments, 376 unique 2-s speech excerpts from the Spoken Wikipedia Corpora^91^ were presented in different background noise conditions and listeners reported the word overlapping the 1-s point by choosing from a closed set of 793 words^16^. The first experiment (44 listeners; 13 female, 30 male, 1 nonbinary; ages 18 to 62 years, median 36) varied the noise level (noiseless and -9, -6, -3, 0, and +3 dB SNR) and included four types of noise: auditory scenes, eight-talker speech babble, instrumental music, and speech-shaped noise spectrally matched to each foreground. The second experiment (47 listeners; 24 female, 23 male; ages 23 to 59 years, median 39) fixed the SNR at -3 dB and included 43 naturalistic auditory textures as noise conditions. At the start of each experiment, listeners completed a headphone check^92^ and level calibration to set a comfortable presentation level, and trials included attention checks. We presented the 25,944 stimuli from both experiments to the model, with foregrounds fixed at 60 dB SPL and the audio resampled from 20 kHz to 50 kHz. Word recognition scores were computed for each of the 69 background noise conditions and compared to those obtained from human listeners.

#### Word and voice recognition with F0-shifted speech

We compared human and model word and voice recognition for F0-shifted speech^26^ using human data from ref.^17^. In an online voice recognition experiment, 112 listeners (46 female, 65 male, 1 nonbinary; ages 20 to 73 years, median 39) heard 4-s excerpts from 37 recognizable public figures. Unbeknownst to the listeners, the F0 on each trial was shifted by -12, -6, -3, 0, +3, +6 or +12 semitones using STRAIGHT^93^. Listeners attempted to identify each voice from a 100-alternative closed set. Voice recognition scores were calculated only for voices that listeners self-reported as familiar in a pre-experiment survey and correctly identified in a subsequent trial block with unshifted clips. Human word recognition scores were measured in a separate online experiment with 22 listeners (8 female, 14 male; ages 25 to 70 years, median 38) in which the same F0 shifts were applied to the clean speech material from the previous word-in-noise experiment. For the model experiment, we applied the same F0 shifts to a set of 373 held-out speech excerpts from talkers familiar to the model (i.e., that were present in the training dataset) and computed word and voice recognition accuracy (across 2611 stimuli) as a function of F0 shift.

#### Environmental sound recognition

We compared human and model environmental sound recognition performance using human data from ref.^25^. In an online experiment 196 listeners (49% female; mean age = 39.9, standard deviation = 13.6 years), listeners heard 2-s auditory scenes comprising 1 to 5 environmental sounds from the GISE-51 evaluation set. Individual sources in the scenes were truncated to 1 s (44.1 kHz; 10-ms half-Hanning onset and offset ramps) and their onset times were sampled uniformly between 0 and 1 s. After each scene, participants were asked whether the scene contained a target environmental sound class chosen from the GISE-51 category labels. Each participant completed 204 trials in which each of the 51 classes were queried 4 times (2 trials for which the correct response was yes and 2 for which the correct response was no). For each of the 51 classes, d’ was calculated as the difference between the z-transformed hit rate and false alarm rate computed across all participants (split half-reliability = 0.97).

To simulate this experiment on our model (which was trained on a single-class environmental sound recognition task), we first trained a new linear readout to perform the same 51-way multi-label detection task. This linear classifier mapped the 512-unit fully connected layer representations of our model’s environmental sound recognition branch to independent probabilities for each of the 51 GISE classes with a sigmoid activation function. The classifier was trained on 240,000 scenes comprising 1 to 5 environmental sounds (each 1 s in duration, with an onset sampled between 0 and 1s and a level between 50 and 70 dB SPL) from the GISE-51 training dataset. The classifier converged in 7500 training steps (batch size 32) using the Adam optimizer with a learning rate of 1e-3 that decreased by a factor of 10 every 2500 steps. We evaluated the classifier on the 19,992 evaluation set scenes from the human experiment, calculated the area under the receiver operating characteristic curve for each of the 51 classes, and converted to d’ to compare with human performance. We note that the model’s d’ values tended to underestimate human performance (Fig. 1i). This is likely because the model representations were optimized for an environmental sound recognition task with a relatively small number of classes and a relatively small dataset of target sounds. Nevertheless, the model captures some elements of human environmental sound recognition behavior, resulting in a higher human-model correlation (r=0.76) than has been reported for traditional models with hand-engineered features (r=0.52) on the same benchmark^25^.

#### Spatial release from masking in different rooms

We compared the model’s spatial speech perception with human data from ref.^27^, who measured speech reception thresholds in 8 normal-hearing listeners (21 to 43 years) for different spatial configurations of speech and noise. Their stimuli comprised syntactically constrained five-word German sentences and speech-shaped noise from the Oldenburg Sentence Test convolved with binaural room impulse responses measured in an anechoic room, an office, and a cafeteria. The speech location was fixed straight ahead, and 50% speech reception thresholds were measured for different rooms and noise azimuths by having listeners repeat each word they recognized. The noise level was adjusted adaptively after each sentence. For the model experiment, we used the same binaural room impulse responses used in the original human experiment but applied them to our English speech materials and middle word recognition task (using the same 2-s stimuli and task described above). Model word recognition accuracy was measured for each room and noise azimuth at SNRs from -24 to +18 dB in 3-dB steps. We fit a sigmoid psychometric function and defined the model’s speech reception threshold as the SNR yielding 50% of the maximum word-recognition performance. We compared spatial release from masking between model and human listeners as the change in speech reception threshold relative to the 0° speech-noise separation, as a function of noise azimuth and room. Absolute speech reception thresholds were not compared because they depend on language and speech materials^94^.

### Model evaluation: psychoacoustic thresholds

We assessed the model’s psychoacoustic sensitivity by training linear classifiers to perform 2I-2AFC discrimination tasks using its internal activations. For each experiment, one or more classifiers were trained on stimuli synthesized to match the corresponding human study as closely as possible. Where necessary, experiments were modified slightly to frame them as 2I-2AFC tasks with 2 s trials (see below). All classifiers were trained on distributions uniformly spanning the relevant stimulus conditions and task difficulties. The test trials were specific points within these same distributions but were sampled independently from the training trials. Training distributions extended well above and below human thresholds, ensuring that classifiers were trained on trials ranging from easy to impossible for human listeners.

#### Linear classifier optimization

Unless otherwise specified, classifiers operated on the ecologically optimized model’s layer 6 activations, corresponding to the output of the final layer normalization operation in the shared trunk. These activations were flattened and passed to a single-unit linear readout with a sigmoid output. Only the weights and bias of this readout were optimized, minimizing the binary cross-entropy loss against ground-truth interval labels (0 = first interval, 1 = second interval). Each classifier was trained for 64,000 trials (2000 steps with batch size 32) using AdamW, with a learning rate of 1e-3 initially, reduced to 1e-4 after 100 steps and 1e-5 after 1000 steps. A weight-decay coefficient of 1e-2 and 50% dropout were used for regularization. This optimization schedule reliably achieved convergence across all experiments.

For analyses of other network layers, including the auditory nerve input representation, the classifier training procedure was identical, although the number of linear weights varied according to the dimensionality of the activations. For layers deeper than the branch point, activations from the four task-specific branches were concatenated and flattened into a single vector before the linear readout. For the untrained network analysis, the same procedure was used, except that the ecologically optimized network weights were replaced by their random initial values. For the psychoacoustically optimized models, the weights of network layers 1-6 were updated jointly with the linear readouts. These models were trained separately for each psychoacoustic task. The network weights were initialized from the ecologically optimized model, and the linear readout was randomly initialized. We used Adam with a learning rate of 1e-5 (matching the ecological training procedure) and fine-tuned the network and readout jointly for 10,000 steps with a batch size of 32 (320,000 trials) for each experiment.

#### Model threshold estimation

Linear classifiers were evaluated on stimulus conditions reported in each study, using the method of constant stimuli and many more trials than the human studies, which typically used adaptive procedures. Separate logistic functions relating the proportion of trials correct to the task difficulty parameter were fit for each condition in each experiment. Unless otherwise noted, model thresholds were defined as the task difficulty yielding 70.7% correct responses. For some stimulus conditions and model variants (particularly the peripheral and untrained models), classifiers never attained 70.7% accuracy. In these cases (in which the threshold was unmeasurably high) we defined the threshold to be the maximum y-axis value of each result plot for the purposes of visualization and statistical analyses.

#### Human threshold extraction

Mean human psychophysical thresholds were manually extracted from figures in published psychoacoustic studies using WebPlotDigitizer (https://apps.automeris.io/wpd4/). Because many studies did not report individual participant thresholds and because our primary aim was to compare group-level central tendencies across many experiments, we did not collect uncertainty estimates within experiments. For the spectrotemporal modulation masking^36^, nonsimultaneous masking^51^, and psychophysical tuning curve^54^ experiments, we used Google Gemini to assist with the initial extraction of data points from figures. The authors subsequently checked all extracted values against the original figures for correctness.

#### Human and model threshold comparisons

We assessed the absolute similarity between human and model thresholds by computing the median model-to-human threshold ratio for each experiment. We assessed the relative similarity across stimulus conditions by computing the Spearman rank correlation coefficient (ρ) between human and model thresholds for each experiment. To quantify the aggregate effects of our model manipulations, we compared median threshold ratios and correlation coefficients between model variants across the 31 psychoacoustic experiments. Statistical significance was assessed with Wilcoxon signed-rank tests and effect sizes were quantified with Cohen’s d and rank-biserial correlation coefficients. When visualizing aggregate distributions of model-to-human threshold ratios (Figs. 3 and 4), the threshold ratios were bootstrapped within experiments and then combined to ensure that each experiment contributed equally to the aggregate distribution. The aggregate distributions of correlation coefficients (Fig. 3) were bootstrapped across experiments.

### Psychoacoustic experiments

All stimuli were generated at 50 kHz. Onset and offset ramps were half-Hanning windows with the specified durations.

#### Pure tone detection in silence

This experiment measured the absolute audibility of pure tones. Training stimuli comprised two 1-s intervals containing either silence or a pure tone (including 50-ms ramps) in random order. The tone’s frequency was sampled log-uniformly between 100 and 20,000 Hz and the level was sampled uniformly between -20 and 100 dB SPL. The tone’s starting phase was randomized. Thresholds were estimated at 125, 250, 500, 750, 1000, 1500, 2000, 3000, 4000, 6000, 8000, 10000, 12000, 14000, and 16000 Hz by evaluating the classifier at sound levels from -20 to 100 dB SPL in 2-dB steps (100 trials per condition). Human thresholds were taken from the ISO 226:2003 standard^95^.

#### Pure tone intensity discrimination

This experiment measured sensitivity to differences in sound intensity. Training stimuli comprised two 500-ms pure tone intervals separated by 500 ms of silence. Both intervals had the same frequency, sampled log-uniformly between 100 and 10,000 Hz, but one interval was presented at a higher intensity than the other (*I* + Δ*I*). The reference level (*I*) was sampled uniformly between 0 and 100 dB SL relative to the ISO 226:2003 standard. The intensity Weber fraction (Δ*I*/*I*) was sampled log-uniformly between 0.01 and 20.0. Thresholds were estimated for 200, 400, 600, 800, 1000, 2000, 4000, and 8000-Hz pure tones at reference levels of 5, 10, 20, 40, and 80 dB SL by evaluating the classifier on 21 log-spaced Weber fractions between 0.0002 and 20 (100 trials per condition). Human thresholds were taken from ref.^46^.

#### Duration discrimination of noise and tone bursts

This experiment measured sensitivity to differences in sound duration. Training stimuli comprised two intervals containing sounds that differed in duration by Δ*t*. The shorter duration t was selected from 13 log-spaced values between 0.156 and 640 ms, and Δ*t* was sampled log-uniformly between 0.1 and 300 ms. Both sounds were centered in their 1-s intervals. There were five stimulus conditions sampled with equal probability: broadband noise (0 to 4 kHz) at 85 or 65 dB SPL, low-pass noise (0 to 1 kHz) at 85 dB SPL, narrowband noise (0.9 to 1.1 kHz) at 85 dB SPL, and a 1-kHz pure tone at 85 dB SPL. Thresholds, defined as the Δ*t* yielding 75% correct responses, were estimated for the same reference durations and stimulus conditions by evaluating the classifier on 31 log-spaced values of Δ*t* between 0.002 and 200 ms (100 trials per condition). Human thresholds were taken from ref.^47^.

#### Temporal gap detection as a function of sound pressure level

This experiment measured the detection of brief silent gaps in broadband noise. Training stimuli comprised two 500-ms broadband (0 to 7 kHz) noise intervals separated by 500 ms of silence. One of the noise intervals included a silent gap with a duration sampled log-uniformly between 1 and 400 ms. Onsets and offsets had 1-ms ramps. Gap durations were calculated from the ramp midpoints (i.e., the smallest possible gap duration was 1 ms). Stimulus levels were sampled uniformly between 10 and 90 dB SPL. Thresholds, defined as the gap duration yielding 73.4% correct responses, were estimated at 20, 30, 40, 50, 60, 70, 80, and 90 dB SPL by evaluating the classifier on 25 log-spaced gap durations between 0.02 ms and 400 ms (100 trials per condition). Human thresholds were taken from ref.^48^.

#### Temporal gap detection as a function of frequency and bandwidth

This experiment measured the detection of brief silent gaps in bandlimited noise. Training stimuli comprised two 400-ms noise intervals separated by 500 ms of silence. One noise interval included a silent gap whose duration was sampled log-uniformly between 0.5 and 250 ms. Onsets and offsets had 0.5-ms ramps and gap durations were calculated from the ramp midpoints. The noise center frequency was sampled log-uniformly between 100 and 10,000 Hz and the bandwidth (expressed as a fraction of the center frequency) was randomly selected from 0.25, 0.375, 0.5, 0.75, and 1. The level of the noise was randomly selected from 10, 25, or 40 dB/Hz SPL. Thresholds were estimated at the same frequencies, bandwidths, and levels by evaluating the classifier on 19 log-spaced gap durations between 0.2 and 200 ms (100 trials per condition). Human thresholds were taken from ref.^49^.

#### Forward masking as a function of frequency and level

This experiment measured the effect of a preceding sound on the detectability of pure tones. Training stimuli comprised two 800-ms intervals separated by 400 ms of silence. At the center of each interval was a 296-ms pure tone masker with frequency sampled log-uniformly between 100 and 5000 Hz and level randomly selected from negative infinity (unmasked condition), 10, 20, 30, 40, 50, 60, 70, 80, or 90 dB SPL. The task was to detect which of the two intervals contained a 24-ms signal pure tone (at the same frequency) that followed the masker with a short delay, sampled log-uniformly between 1 and 100 ms from masker offset to signal onset. All onsets and offsets had 12-ms ramps. The signal level was sampled uniformly between -20 and 100 dB SPL. Detection thresholds were estimated for different combinations of frequency (125, 250, 500, 1000, 4000 Hz), masker level (unmasked, 20, 40, 50, 60, 70, 80, and 90 dB SPL), and masker-to-signal delay (4, 8, 16, and 32 ms) by evaluating the classifier on signal levels from 0 to 120 dB SPL in 5-dB steps (50 trials per condition). Masking in dB was calculated as the difference between masked and unmasked thresholds. To prevent infinite masking values in the deterministic model, in which absolute audibility is limited only by numerical precision, unmasked thresholds were lower bounded by the ISO 226:2003 hearing thresholds. Human masking values were taken from ref.^50^.

#### Forward, backward, and combined nonsimultaneous masking

This experiment measured the effect of preceding and following sounds on the detectability of a brief tone. Training stimuli comprised two 650-ms intervals separated by 500 ms of silence. Both intervals contained identical broadband masking noise (0 to 9 kHz, presented at 40 dB/Hz SPL) but only one contained the signal (a 2-ms 6-kHz pure tone pip including 1-ms ramps). We estimated thresholds for seven conditions: no masking, simultaneous masking (signal centered in a 400-ms masker), forward masking alone (signal placed after a 200-ms masker), backward masking alone (signal placed before a 200-ms masker), combined forward and backward masking (signal placed between two 200-ms maskers) with symmetric delays (*t*_1_ = *t*_2_), and two combined masking conditions with asymmetric delays. In the two asymmetric conditions, either the pre- or post-signal delay was increased by 20 ms (*t*_1_ = *t*_2_ + 20 ms or *t*_2_ = *t*_1_ + 20 ms). The smaller delay was sampled uniformly between -11 and 100 ms, with negative delays indicating that the signal overlapped the masker. The signal level was sampled uniformly between 0 and 120 dB SPL. Thresholds were estimated at 14 delays between 0 and 100 ms by evaluating the classifier on signal levels from 0 to 120 dB SPL in 5-dB steps (50 trials per condition). Human thresholds were taken from ref.^51^. To simplify presentation, thresholds for the asymmetric masking conditions are not shown in Fig. 7g but were included in all summary metrics.

#### Forward masking with broadband noise

This experiment measured the effect of preceding noise bursts on the detectability of a brief tone. Training stimuli comprised two 800-ms intervals separated by 400 ms of silence. Centered in each interval was a 200-ms broadband (0 to 8 kHz) noise masker with level sampled uniformly between 10 and 90 dB SPL. Only one interval contained the signal, a 10-ms 4-kHz pure tone pip including 5-ms ramps. The signal level was sampled uniformly between -20 and 100 dB SPL and the signal onset was delayed relative to the masker offset. Masker-to-signal delays were sampled uniformly between -50 and 200 ms. Thresholds were estimated at masker levels of 40, 60 and 80 dB SPL and 13 delays between -20 and 150 ms by evaluating the classifier on signal levels from -20 to 100 dB SPL in 5-dB steps (50 trials per condition). Human thresholds were taken from ref.^52^.

#### Forward masking with on- and off-frequency pure tones

This experiment measured the effect of preceding pure tones on the detectability of a brief tone. Training stimuli were identical to those in the previous experiment except the broadband noise maskers were replaced with either on-frequency (4 kHz, matching the signal frequency) or off-frequency (2.4 kHz) pure tones. Human thresholds were taken from ref.^52^, which repeated the experiment of ref.^59^. Thresholds were estimated for both maskers at 9 levels between 10 and 90 dB SPL and at delays of 0, 10, and 30 ms by evaluating the classifier on signal levels from -20 to 100 dB SPL in 5-dB steps (50 trials per condition).

#### Critical band masking patterns

This experiment measured the effect of concurrent noise on the detectability of pure tones. Training stimuli comprised two 1-s intervals of bandlimited masking noise presented at 31 dB/Hz SPL with up to ±5 dB of level roving. The low frequency cutoff and bandwidth of the noise were sampled log-uniformly from 400 to 10,000 Hz and 200 to 4000 Hz, respectively. The task was to detect a 1-s pure tone (including 25-ms ramps) added to one of the intervals. The tone frequency was sampled log-uniformly from 500 to 8000 Hz. For evaluation, the masker level (31 dB/Hz SPL) and low frequency cutoff (3030 Hz) were fixed. Signal detection thresholds were estimated for masker bandwidths of 60, 84, 104, 175, 288, 470, 720, and 1220 Hz and 21 log-spaced tone frequencies between 2400 and 4800 Hz by evaluating the classifier on signal levels from 0 to 80 dB SPL in 2-dB steps (50 trials per condition). Human thresholds were taken from ref.^53^.

#### Psychophysical tuning curves

This experiment measured the level of concurrent narrowband noise required to mask pure tones at different frequencies. Training stimuli comprised two 220-ms narrowband (25-Hz wide) noise intervals separated by 500 ms. The signal to detect was a 20-ms pure tone pip (including 10-ms ramps) added to one of the maskers. The signal frequency was either 500, 1000, 2000, or 4000 Hz presented at a fixed 43, 52, 54.5, or 53 dB SPL, respectively, to match the mean of two human psychophysical tuning curves from ref.^54^. The narrowband masker level was sampled uniformly between 0 and 120 dB SPL. Continuous notched noise was added to all trials to reduce off-frequency listening. For each signal frequency (*f*) the noise spanned 0.6 ⋅ *f* to 1.4 ⋅ *f* with a notch between 0.925 ⋅ *f* and 1.075 ⋅ *f*. Notched-noise levels were -7.5, -0.5, 5, and 1 dB/Hz SPL for the four frequencies. These levels were 10 dB lower than those used for the human listeners, which was necessary to measure tuning curves from our peripheral and untrained model variants. Tuning curves for the ecologically and psychoacoustically optimized models were unaffected by this modification. Classifiers were evaluated on narrowband masker levels from 0 to 120 dB SPL in 5-dB steps (100 trials per condition). For statistical analyses, human and model tuning curves at each frequency were converted to dB SNR by subtracting the threshold masker level from the signal level, ensuring that lower thresholds always corresponded to better performance.

#### Sinusoidal and narrowband masking patterns

This experiment measured how simultaneous masking varied with signal frequency and masker level. Separate linear classifiers were trained to detect either pure tones or narrowband (80-Hz wide) noise signals. Both signals were masked by either 1-kHz pure tones or narrowband (940 to 1040 Hz) noise. Training stimuli comprised two 220-ms masker intervals separated by 500 ms of silence. The 220-ms (including 10-ms ramps) signal was added to one of the maskers. Masker levels were sampled from no masker (unmasked condition), 45, 65, and 85 dB SPL. Signal frequency was sampled log-uniformly between 125 and 8000 Hz and signal level was sampled uniformly between 0 and 100 dB SPL. For tone-on-tone masking trials, the signal and masker tone always started 90° out of phase. Thresholds were estimated for all signal-masker pairings, at each masker level, and at 11 signal frequencies between 250 and 4000 Hz by evaluating the classifier on signal levels from -20 to 120 dB SPL in 5-dB steps (50 trials per condition). Human thresholds were taken from ref.^55^.

#### Pure tone frequency discrimination

Sensitivity to frequency differences was measured as a function of pure tone frequency with three different paradigms: frequency change detection, up/down frequency discrimination, and frequency modulation detection. We trained 21 separate classifiers to perform each task at seven reference frequencies: 250, 500, 1000, 2000, 4000, 6000, and 8000 Hz. For the change detection task, training stimuli comprised two 530-ms intervals separated by 300 ms. Each interval contained two 240-ms pure tones (including 20-ms ramps) separated by 50 ms. In one interval both tones had the same frequency (*f*). In the other interval, the tones had different frequencies (*f* and *f* + Δ*f*). The task was to report which interval included the frequency change. For the up/down discrimination task, two intervals each contained a single 240-ms pure tone (including 20-ms ramps), one with frequency f and the other with frequency *f* + Δ*f*. The task was to report which interval had the higher frequency. For the modulation detection task, one interval contained a 1-s pure tone, and the other contained a 1-s frequency-modulated tone (including 50-ms ramps). The frequency of the modulated tone varied sinusoidally around the pure tone frequency (*f*) with a peak-to-peak amplitude of Δ*f* and a modulation rate of 2, 5, or 10 Hz. To prevent classifiers from exploiting low-level artifacts from pure tone frequency modulation, trials were band-pass filtered and included continuous Gaussian noise (0 dB/Hz SPL). For all three paradigms, *f* was sampled uniformly within ±0.5 octaves of the reference frequency and the frequency Weber fraction Δ*f*/*f* was sampled log-uniformly between 5e-4 and 5e-1. Training trials also included ±5 dB of level roving across intervals. Thresholds for each task were estimated at each reference frequency by evaluating the classifier on 43 values of Δ*f*/*f* log-spaced between 5e-8 and 5e-1 (100 trials per condition). Human thresholds for all three experiments were taken from ref.^57^.

#### Formant frequency discrimination

This experiment measured sensitivity to differences in the spectral envelopes of vowel sounds. Training stimuli comprised two 160-ms intervals (including 5-ms ramps) separated by 160 ms of silence. One interval contained a synthetic vowel, and the other contained the same vowel with the first or second formant frequency shifted up or down. Ten distinct vowels were synthesized using a source-filter model with standard formant frequencies and bandwidths (following ref.^56^). Vowel F0s were sampled log-uniformly between 100 and 400 Hz. Four classifiers were trained separately to detect upward or downward shifts in either the first or second formant. During training, frequency shifts, quantified as Weber fractions (Δ*f*/*f*) relative to the standard formant frequency, were sampled log-uniformly between 5e-3 and 5e-1. Thresholds for each vowel, formant, and direction were estimated by evaluating the classifiers on 25 log-spaced values of Δ*f*/*f* between 5e-5 and 5e-1 (100 trials per condition). Thresholds were averaged across the two shift directions and compared to human data from ref.^56^.

#### F0 discrimination

This experiment measured sensitivity to differences in the F0 of complex tones. Training stimuli comprised two 500-ms harmonic complex tone intervals (including 30-ms ramps) separated by 500 ms. The two complex tones had the same spectral envelope but different F0s (*F*0*_ref_* and *F*0*_ref_* + Δ*F*0). The task was to report which interval had the higher F0. For a given trial, there were two possible spectral envelopes: a low spectral region (imparted by a 4^th^-order band-pass filter with a -3 dB passband of 2.5 to 3.5 kHz) and a high spectral region (−3 dB passband of 5 to 7 kHz). For low spectral region trials, *F*0*_ref_* was sampled from 50, 75, 100, 150, 200, and 300 Hz. For high spectral region trials, *F*0*_ref_* was sampled from 100, 150, 200, 300, 400, and 600 Hz. All trials were presented in continuous noise with a flat 15 dB/Hz SPL spectrum level below 600 Hz and a 2-dB/octave roll-off above 600 Hz. Individual harmonic components were scaled to 48.3 dB SPL (15 dB SL for human listeners) before filtering. The nominal lowest harmonic number in each complex was defined as the first harmonic attenuated less than 15 dB by the band-pass filter. Harmonics were combined in either sine or random phase on each trial. During training, the Weber fraction Δ*F*0/*F*0*_ref_* was sampled log-uniformly between 1e-5 and 1. Thresholds were estimated for each spectral region, *F*0*_ref_*, and harmonic phase condition by evaluating the classifier on 37 log-spaced Δ*F*0/*F*0*_ref_*values between 1e-6 and 1 (100 trials per condition). Human thresholds were taken from ref.^58^.

#### Amplitude modulation detection

These experiments measured sensitivity to fluctuations in sound amplitude across time. Training stimuli comprised two 1-s observation intervals, each including 200-ms ramps. Both intervals contained the same noise or tone carrier, and the task was to detect a sinusoidal amplitude modulation imposed on one of the intervals. Separate classifiers were trained for eight different carriers across two experiments. The first experiment examined the effect of carrier bandwidth, comparing broadband noise (0 to 6 kHz), a 5-kHz pure tone, and three narrowband noise conditions (3, 31, and 314-Hz wide, each centered at 5 kHz). The second experiment fixed the carrier bandwidth (50-Hz wide, centered at 5 kHz) and examined the effect of carrier envelope statistics, comparing Gaussian noise, multiplied noise, and low-noise noise conditions. Multiplied noise was generated by multiplying a 5 kHz sinusoid with a low-pass (0 to 25 Hz) Gaussian noise modulator. Low-noise noise was generated from bandlimited Gaussian noise by repeatedly dividing the waveform by its Hilbert envelope and re-imposing its original 50-Hz bandwidth in the frequency domain (10 iterations). For the broadband noise condition, stimuli were presented at 77 dB SPL, and the modulation rate was sampled log-uniformly between 2 and 2000 Hz. For the seven other conditions, stimuli were presented at 65 dB SPL, and the modulation rate was sampled log-uniformly between 2 and 200 Hz. During training, modulation depths were sampled uniformly between -30 and 0 dB. Thresholds were estimated for each carrier as a function of modulation rate by evaluating the classifiers on modulation depths from -40 to 0 dB in 1-dB steps (100 trials per condition). Human thresholds for the bandwidth experiment were taken from ref.^29^ (broadband condition) and ref.^4^ (narrowband conditions). Human thresholds for the noise type experiment were taken from ref.^31^.

#### Amplitude modulation masking

This experiment measured the masking of sinusoidal amplitude modulation by concurrent modulation. Training stimuli comprised two 600-ms observation intervals separated by 500 ms of silence. Both intervals contained broadband (1 to 4 kHz) noise carriers sinusoidally amplitude modulated at the same masker modulation rate. The task was to detect an additional target modulation imposed on one of the intervals. Separate classifiers were trained to detect target modulation rates of 4, 16, 64, and 256 Hz. The masker modulation rate was sampled uniformly within ±2 octaves of the target modulation rate. The masker modulation depth was fixed at -10 dB, and the target modulation depth was sampled uniformly between -30 and 0 dB. Thresholds for each target modulation were estimated as a function of masker modulation rate by evaluating the classifiers on target modulation depths from -40 to 0 dB in 1-dB steps (100 steps per condition). Human thresholds were taken from ref.^32^.

#### Spectral modulation detection

This experiment measured sensitivity to fluctuations in sound amplitude across the frequency dimension. Training stimuli comprised two 400-ms bandlimited noise intervals (including 10-ms ramps) separated by 400 ms of silence. The noise in both intervals had the same frequency band, sampled from 200-400, 400-800, 800-1600, 1600-3200, 3200-6400, 6400-12800, 200-800, 800-3200, 3200-12800, 200-1600, 1600-12800, and 200-12800 Hz. One interval had a flat spectral envelope presented at 35 dB/Hz SPL. The other had a sinusoidal spectral envelope with a modulation rate sampled from 1/4, 1/3, 1/2, 1, 4/3, 3/2, 2, 3, 4, 5, 7, and 10 cycles per octave, a peak-to-valley modulation depth sampled uniformly between 0 and 20 dB, and a random starting phase. Modulation rate sampling was constrained such that the noise bandwidth contained at least 1.5 modulation cycles. Presentation levels within trials were roved by up to ±5 dB. Modulation detection thresholds were estimated for all combinations of noise band and spectral modulation rate by evaluating the classifier on modulation depths from 0 to 20 dB in 1-dB steps (100 trials per condition). Human thresholds were taken from ref.^33^.

#### Spectral modulation masking

This experiment measured the masking of spectral modulation by concurrent modulation. Stimuli were similar to the previous experiment, except an additional “masker” spectral modulation was added to both intervals. The masker spectral modulation had a fixed depth of 15 dB and a rate that was chosen at random from 0 (unmasked condition), 1, 3, or 5 cycles per octave. The target spectral modulation rate was sampled log-uniformly between 0.2 and 20 cycles per octave with a depth sampled uniformly between 0 and 25 dB. Thresholds were estimated as a function of target modulation frequency for three noise bands (200-12800, 800-1600, and 6400-12800 Hz) and four masker modulation rates by evaluating the classifier on target modulation depths of 0 to 25 dB in 1-dB steps (100 trials per condition). Human thresholds were taken from ref.^35^.

#### Spectrotemporal modulation detection

This experiment measured sensitivity to fluctuations in sound amplitude across time and frequency. Training stimuli comprised two 1-s observation intervals, each including 10-ms ramps. One interval contained a broadband noise carrier constructed by summing 1000 random-phase pure tones with frequencies log-spaced from 140 to 7,340 Hz. The other interval contained the same carrier with a sinusoidal spectrotemporal modulation imposed. The spectrotemporal modulation was parametrized with a spectral modulation rate (sampled from 0.25, 0.5, 1, 2, 4, and 8 cycles per octave), a temporal modulation rate (sampled from ±1, ±2, ±4, ±8, ±16, ±32, ±64, and ±128 Hz), and a modulation depth (sampled uniformly between -30 and 0 dB, with 0 dB corresponding to 100% modulation). The sign of the temporal modulation rate determined the orientation of the spectrotemporal ripple, with negative rates corresponding to downward moving ripples. Presentation levels were sampled independently for the two intervals between 60 and 80 dB SPL. Modulation detection thresholds were estimated for all combinations of spectral and temporal modulation rates by evaluating the classifier on modulation depths from -40 to 0 dB in 2-dB steps (100 trials per condition). Human thresholds were taken from ref.^34^. The stimuli in the human experiment were constructed using only 92 tones. We increased this number to 1000 to prevent classifiers from exploiting spectral aliasing artifacts^96^.

#### Spectrotemporal modulation masking

This experiment measured the masking of spectrotemporal modulation by concurrent modulation. Training stimuli comprised two 500-ms observation intervals (including 25-ms ramps) separated by 500 ms of silence. Both intervals contained broadband noise (constructed by summing 541 random-phase pure tones with frequencies between 8 and 5200 Hz) with the same masker spectrotemporal modulation imposed. The task was to detect an additional target spectrotemporal modulation imposed on one of the intervals. Separate classifiers were trained to detect four target modulations with respective spectral and temporal modulation rates (in cycles per octave, Hz) of (1, -8), (1, +8), (0.5, +4), and (1, +16). Target modulation depth ranged from -30 to 0 dB. Masker spectral and temporal modulation rates were randomly sampled for each trial. For each target modulation, thresholds were estimated for 25 combinations of masker spectral and temporal modulation rates surrounding the target by evaluating the classifier on modulation depths from -40 to 0 dB in 2-dB steps (100 trials per condition). Human thresholds were taken from ref.^36^.

#### Interaural correlation discrimination

This experiment measured sensitivity to the correlation between sound waveforms at the two ears. Training stimuli comprised two 300-ms intervals (including 15-ms ramps) separated by 100 ms of silence. Both intervals contained noise presented at 75 dB SPL with a bandwidth sampled log-uniformly between 1 and 10,000 Hz. For bandwidths below 1000 Hz, the noise was centered at 500 Hz. For larger bandwidths, the noise was low-pass. In one interval, the noise was generated independently for the left and right ear, producing a reference interaural correlation of 0. In the other interval, the interaural correlation was sampled uniformly between 0 and 1. Given independent noise waveforms *x*_1_ and *x*_2_, an interaural correlation *c* was achieved by presenting *x*_1_ to the left ear and 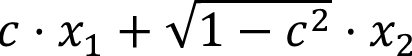 to the right ear. The task was to report which interval had the higher interaural correlation. A separate classifier was trained with a reference interaural correlation of 1. In this case, the task was to report which interval had the lower interaural correlation. Thresholds were estimated for bandwidths of 3, 10, 40, 115, 1000, and 4500 Hz by evaluating the classifiers on 101 log-spaced interaural correlation differences between 1e-5 and 1 (100 trials per condition). Human thresholds were taken from ref.^40^.

#### Interaural level difference discrimination

This experiment measured sensitivity to level differences between the two ears. Training stimuli comprised two 500-ms pure tone intervals (including 15-ms ramps) separated by 500 ms of silence. The frequency on each trial was sampled log-uniformly between 100 and 10,000 Hz. Stimuli were presented at 70 dB SPL with up to ±5 dB level roving across intervals. Each trial was parametrized by two interaural level differences (ILDs): a reference (*I*), sampled uniformly from 0 to 15 dB, and a delta (Δ*I*), sampled uniformly from ±15 dB. The tones in the two intervals had ILDs equal to *I* ± (Δ*I*/2). The task was to report which interval had the larger ILD, with positive ILDs favoring the right ear. Thresholds were estimated at five frequencies (200, 500, 1000, 2000, and 5000 Hz) and three reference ILDs (0, 9, and 15 dB) by evaluating the classifier on 25 log-spaced Δ*I* magnitudes between 0.01 and 10 dB (100 trials per condition). Human thresholds were taken from ref.^37^.

#### Interaural time difference discrimination

This experiment measured sensitivity to time delays between the two ears. Training stimuli comprised two 500-ms pure tone intervals (including 100-ms ramps) presented at 70 dB SPL and separated by 400 ms of silence. The frequency on each trial was sampled log-uniformly between 200 and 2000 Hz. The two intervals had interaural phase differences (IPDs) of ±Δ*IPD*/2, where Δ*IPD* was sampled uniformly between 0 and 2π. The task was to report which interval had the positive right-leading IPD. Thresholds were estimated at frequencies of 250, 500, 700, 800, 900, 1000, 1200, 1250, 1300, 1350, and 1400 Hz by evaluating the classifier on 49 log-spaced Δ*IPD*s between 0.0001 and 1 (100 trials per condition). Threshold Δ*IPD*s were converted to Δ*ITD*s and compared with human thresholds from ref.^38^.

#### Envelope interaural time difference discrimination

This experiment measured sensitivity to time delays between the envelopes of sounds presented to the two ears. Training stimuli comprised two 300-ms tone intervals (including 20-ms ramps) and separated by 400 ms of silence. Stimuli were presented at 75 dB SPL. Each trial contained either pure tones, sinusoidally amplitude-modulated (SAM) tones, or transposed tones. For SAM and transposed tones, carrier frequencies were sampled log-uniformly between 2828 and 14,142 Hz and envelope frequencies were sampled log-uniformly between 23 and 1448 Hz. Pure tone frequencies were sampled from the same range as the envelope frequencies. One interval contained no IPD, whereas the other contained an IPD sampled uniformly between 0 and π. For SAM and transposed tones, the IPD was imposed only on the temporal envelope. For pure tones, the IPD was imposed on the fine structure. IPD detection thresholds were estimated for SAM and transposed tones with carrier frequencies of 4, 6, and 10 kHz and envelope / pure tone frequencies of 32, 64, 128, 256, 512, and 1024 Hz by evaluating the classifier on 49 log-spaced IPDs between 0.0001 and 1 (100 trials per condition). Threshold IPDs were converted to ITDs and compared to human thresholds from ref.^39^. The human study did not measure thresholds at 1024 Hz; however, because models exhibited envelope ITD sensitivity at high frequencies, we included this condition and extrapolated human pure tone thresholds from ref.^38^. SAM and transposed-tone thresholds were assumed to remain unmeasurably high.

#### Minimum audible angles

This experiment measured sensitivity to differences in the horizontal direction from which sounds were presented. Training stimuli comprised two successive 1-s pure tones, each including 70-ms ramps. The tones had the same frequency, sampled log-uniformly between 200 and 12,500 Hz, but were presented at different azimuths in the right frontal hemifield. One tone was presented at a reference azimuth (*θ*), randomly chosen from 0, 30, 45, 60, and 75°, and the other at *θ* + Δ*θ*, with Δ*θ* sampled log-uniformly between 0.5° and 50° subject to *θ* + Δ*θ* ≤ 90°. The task was to report which tone was farther to the right. Tones were presented at 50 dB SL and spatialized using linearly interpolated KEMAR HRTFs, measured every 5° in azimuth at 0° elevation. Thresholds were estimated at 13 frequencies between 250 and 10,000 Hz for each reference azimuth by evaluating the classifier on 31 log-spaced Δ*θ* values between 0.05 and 0.5 (100 trials per condition). Human thresholds were taken from ref.^41^.

#### Binaural masking level differences

Four experiments measured the effects of interaural phase on the detectability of tones in noise. In each experiment, training stimuli comprised two intervals containing masking noise (*N*), one of which also contained a signal tone (*S*).

The first experiment measured signal-in-noise detection thresholds as a function of frequency and interaural phase. The signal was a 1-s pure tone, including 30-ms ramps, with frequency sampled log-uniformly between 80 and 6000 Hz. The masker was broadband noise spanning 0 to 7 kHz and presented at 59.1 dB/Hz SPL. Signal level was sampled uniformly between 50 and 100 dB SPL. On each trial, masking noise was presented either identically to the left and right ear (*N*_0_) or 180° out of phase between the ears (*N_π_*). The signal was presented identically to the two ears (*S*_0_), 180° out of phase (*S_π_*), or monaurally to a random ear (*S_M_*). Thresholds were estimated for all six pairings of signal and masker phase at signal frequencies of 100, 200, 500, 1000, 2000, and 5000 Hz by evaluating the classifier on signal levels from 40 to 110 dB SPL in 2.5-dB steps (50 trials per condition). Human thresholds were taken from ref.^42^.

The second experiment measured signal-in-noise detection thresholds for pure tones as a function of interaural phase, signal frequency, and masker bandwidth. Signal frequency was sampled log-uniformly between 100 and 10,000 Hz. The masker was centered at the signal frequency with a bandwidth sampled log-uniformly between 1 and 10,000 Hz. A 350-ms signal (including 50-ms ramps) was centered in one of two 450-ms masker intervals at an SNR sampled uniformly between -30 and +15 dB. Both intervals were presented at 70 dB SPL. *N*_0_*S*_0_, *N_π_S*_0_, and *N*_0_*S_π_* thresholds were estimated for signal frequencies of 125, 250, 500, 1000, 2000, and 4000 Hz and masker bandwidths of 5, 10, 25, 50, 100, 250, 500, 1000, 2000, 4000, 8000 Hz, subject to the constraint that masker bandwidth did not exceed twice the signal frequency, by evaluating the classifier on SNRs from -45 to +15 dB in 3-dB steps (50 trials per condition). Human thresholds were taken from ref.^43^.

The third experiment measured signal-in-noise detection thresholds for pure and transposed tones as a function of interaural phase and masker bandwidth. The five possible signals were pure tones at 125, 250, or 4000 Hz and transposed tones with 125- or 250-Hz envelopes imposed on a 4000-Hz carrier. Masker bandwidth was sampled log-uniformly between 1 and 1000 Hz, centered at the signal frequency. Durations, SNRs, and presentation levels matched those of the previous experiment. *N*_0_*S*_0_, *N_π_S*_0_, and *N*_0_*S_π_* thresholds were estimated for each signal at masker bandwidths of 5, 10, 25, 50, 100, 250, and 500 Hz by evaluating the classifier on SNRs from -45 to +15 dB in 3-dB steps (50 trials per condition). Human thresholds were taken from ref.^44^.

The fourth experiment measured signal-in-noise detection thresholds for a 500-Hz pure tone as a function of masker interaural correlation and signal interaural phase. The signal duration was 150 ms, including 25-ms ramps and the signal level was sampled uniformly between 40 and 100 dB SPL. The masking noise spanned 100 to 3000 Hz and was presented at 50 dB/Hz SPL. Thresholds were estimated for *S*_0_ and *S_π_* signals in masking noise with interaural correlations of 0, ±0.25, ±0.5, ±0.65, ±0.8, ±0.9, and ±1 by evaluating the classifier on signal levels from 40 to 110 dB SPL in 2.5-dB steps (100 trials per condition). Human thresholds were taken from ref.^45^.

### Error bars

Error bars in Fig. 1 and Supplementary Fig. 1 indicate ±1 standard error of the mean. For human results, standard errors were computed across participants. For model results, standard errors were computed by bootstrapping across trials. Because the model was evaluated on large numbers of trials, the model standard errors were low, often much smaller than the plot markers. For this reason, model error bars are not shown for individual psychoacoustic thresholds (Figs. 5-8). Human error bars are not shown because individual participant data was not available for most of the psychoacoustic experiments (because the data were sourced from figures in the original papers).

## ACKNOWLEDGMENTS

We thank Rainer Beutelmann for sharing the BRIRs for the spatial release from masking experiment and members of the McDermott lab for their helpful feedback on the manuscript. This work was supported by the William Demant Foundation (grant 24-5089 to M.R.S.) and the National Institutes of Health (grant R01DC021464 to J.H.M.).

## SUPPLEMENTARY INFORMATION

**Supplementary Figure 1.**
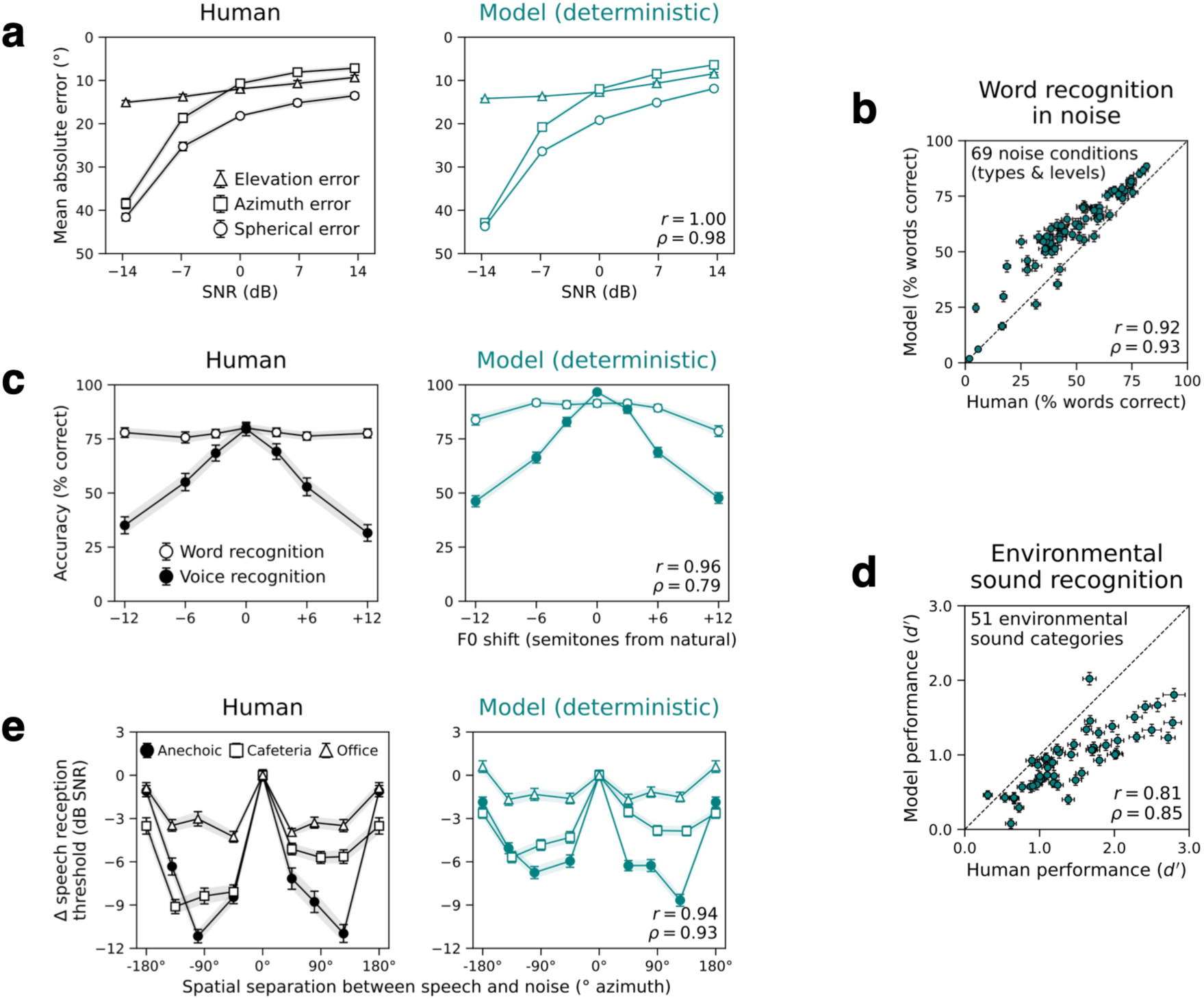
The fully deterministic model optimized for ecological hearing tasks exhibits human-level sound localization and recognition task performance. (**a**) Comparison of human and model localization performance, assessed by measuring mean absolute azimuth, elevation, and spherical error as a function of signal-to-noise ratio. (**b**) Comparison of human and model word recognition accuracy across 69 background noise conditions. (**c**) Human and model word and voice recognition accuracy measured using speech from familiar talkers whose voice has been F0-shifted. (**d**) Comparison of human and model environmental sound recognition performance across 51 different sound classes. (**e**) Human and model speech reception thresholds, measured as a function of the spatial separation between speech and noise in three acoustically distinct rooms. Human-model similarity for each task is quantified using Pearson (r) and Spearman (ρ) correlation coefficients. Human data in (**a**), (**b**), and (**c**) are from ref.^17^. Human data in (**d**) and (**e**) are from refs.^25,27^, respectively. Error bars indicate ±1 s.e.m.

**Supplementary Figure 2.**
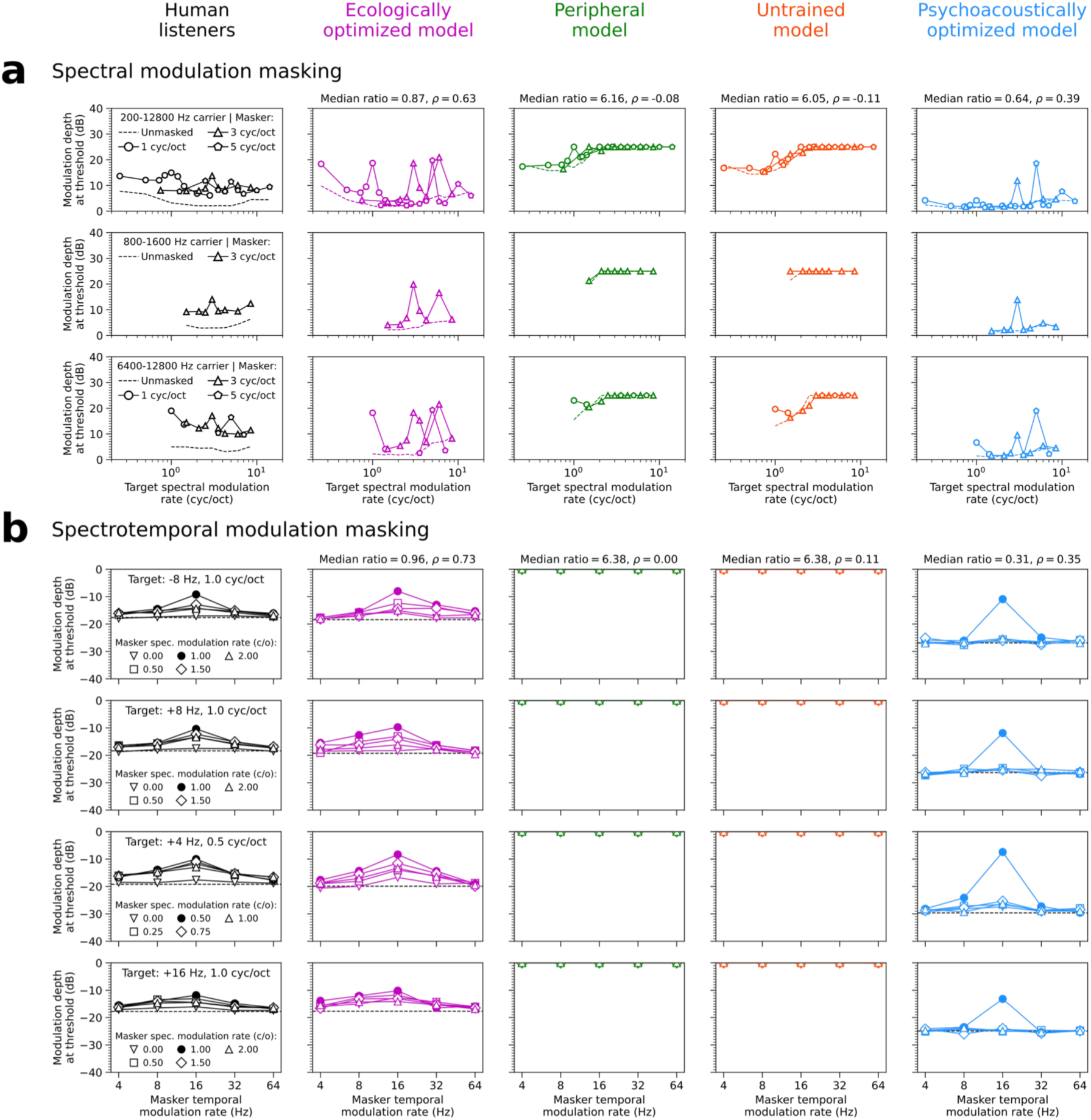
Additional modulation masking experiments. (**a**) Spectral modulation detection thresholds measured in the presence of different masker spectral modulations (different lines) imposed on different carrier noise bands (different rows). (**b**) Spectrotemporal modulation detection thresholds measured for different combinations of target and masker modulations. Rows correspond to four different target spectrotemporal modulations. Masker spectrotemporal modulations are specified along the x-axis (temporal modulation rates) and the different lines (spectral modulation rates).

**Supplementary Figure 3.**
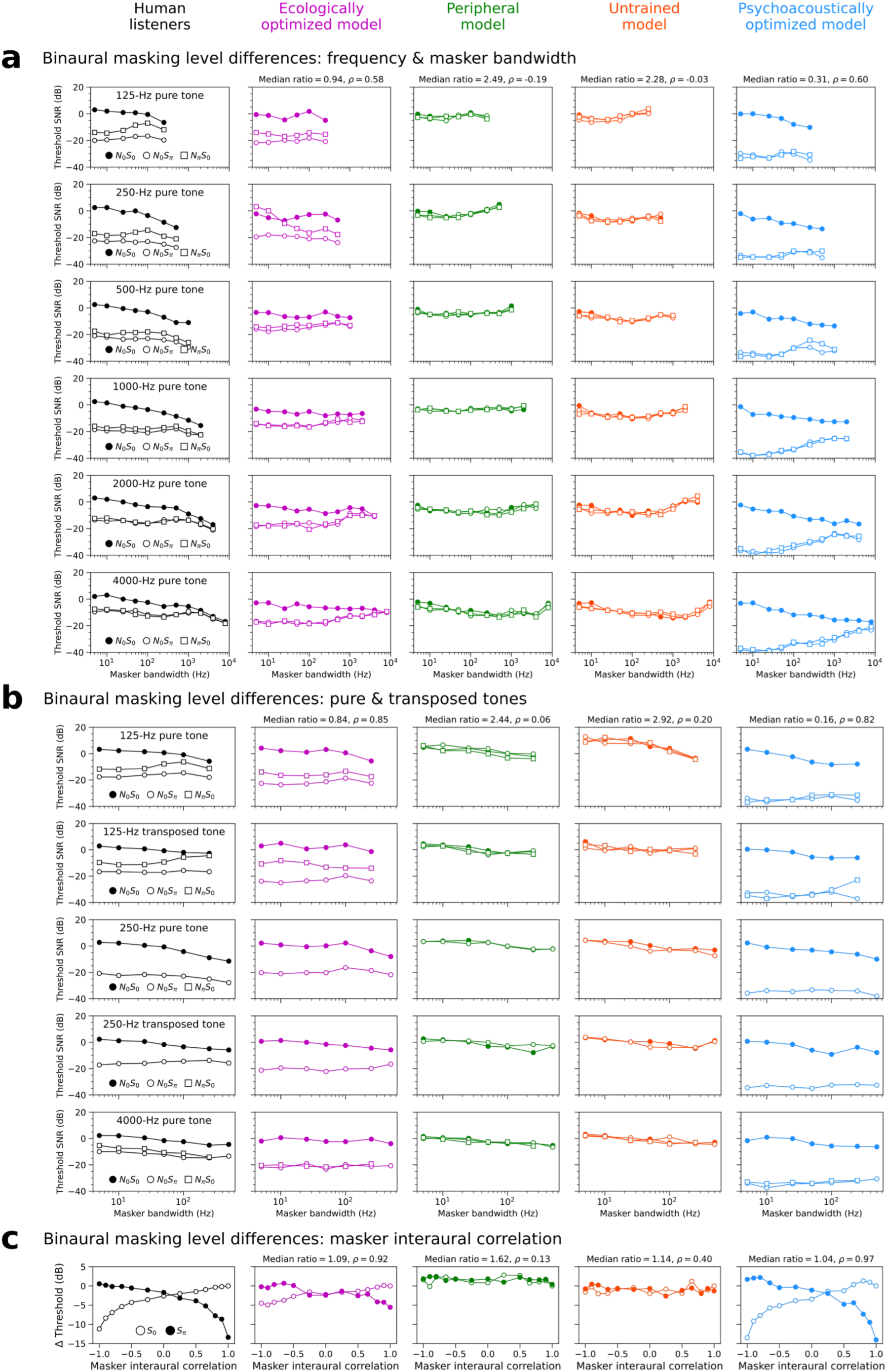
Additional binaural masking level difference experiments. (**a**) Binaural tone-in-noise detection thresholds measured as a function of pure tone frequency, masker bandwidth, and interaural phase condition. The interaural phase of the noise (N) and signal (S) are each denoted with subscripts: 0 = in phase between the ears, π = 180° out of phase. (**b**) Analogous thresholds measured for pure tones and transposed tones with 4000 Hz carrier frequencies. (**c**) Binaural masking level differences (BMLDs) measured as a function of masker interaural correlation. BMLDs indicate the change in detection threshold relative to the *S*_0_ condition with a masker interaural correlation of one. Median model-to-human threshold ratios and human-model Spearman correlation coefficients (ρ) are computed across the stimulus conditions within each experiment.

**Supplementary Figure 4.**
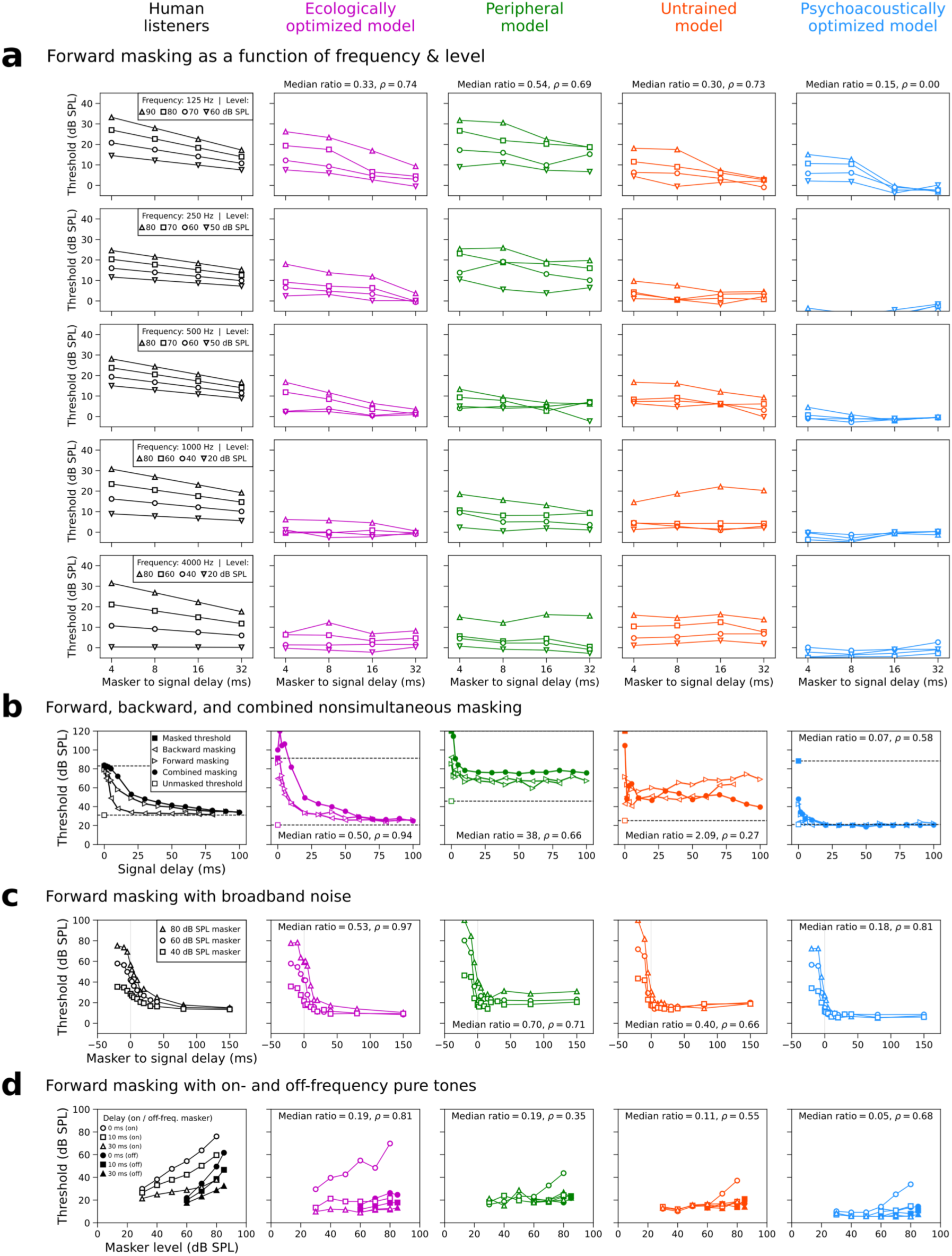
Nonsimultaneous masking experiments. (**a**) Forward-masked pure tone detection thresholds measured as a function of frequency (different rows), masker levels (different lines), and the temporal separation between the masker offset and the signal onset (x-axis). (**b**) Detection thresholds for a 6000-Hz pure tone pip as a function of the temporal separation between signal and masker. In the backward masking condition, the masker occurs after the signal. In the forward masking condition, the masker occurs before the signal. In the combined masking condition, maskers occur both before and after the signal. (**c**) Forward-masked detection thresholds for a 4000-Hz pure tone as a function of masker-to-signal delay for three different levels of broadband noise masker. (**d**) Forward-masked detection thresholds for a 4000-Hz pure tone as a function of masker level and delay for on- and off-frequency pure tone maskers (4000 and 2400 Hz, respectively).

**Supplementary Figure 5.**
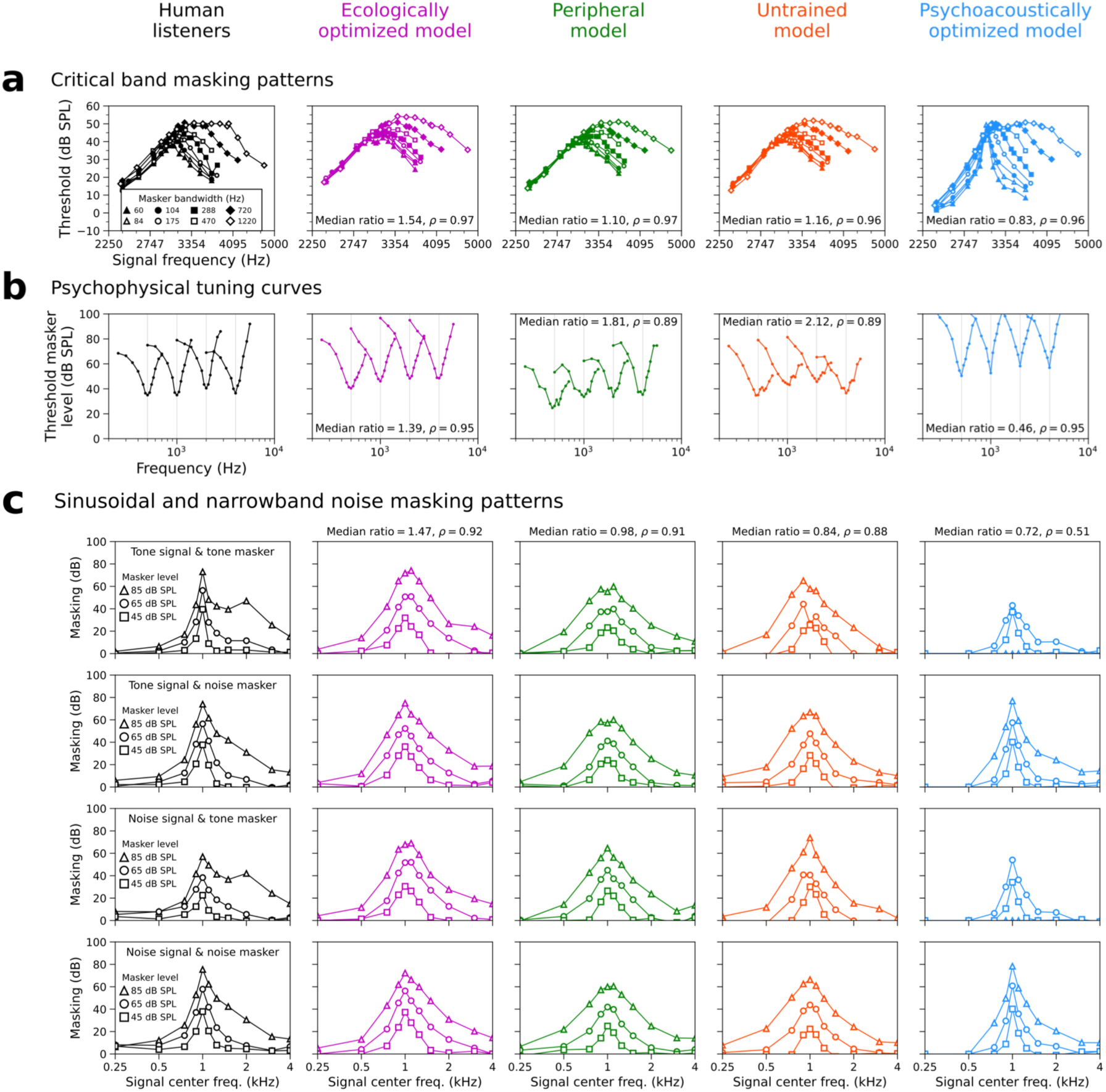
Simultaneous masking experiments. (a) Pure tone detection thresholds measured in the presence of masking noise whose low frequency cutoff was fixed at 3030 Hz and whose bandwidth varied. (b) Psychophysical tuning curves show the level of narrowband noise needed mask a pure tone with frequency 500, 1000, 2000, or 4000 Hz. The narrowband noise was 25-Hz wide and centered at the frequency plotted on the x-axis. (c) Sinusoidal and narrowband noise masking patterns. In this masked detection experiment, the signal and the masker were each either a pure tone or an 80-Hz wide narrowband noise. The four possible signal-masker pairings are shown in the four rows. The masker was always centered at 1 kHz and signal detection thresholds were measured as a function of signal center frequency for three masker levels. The amount of masking was computed as the threshold increase relative to unmasked detection.

**Supplementary Figure 6.**
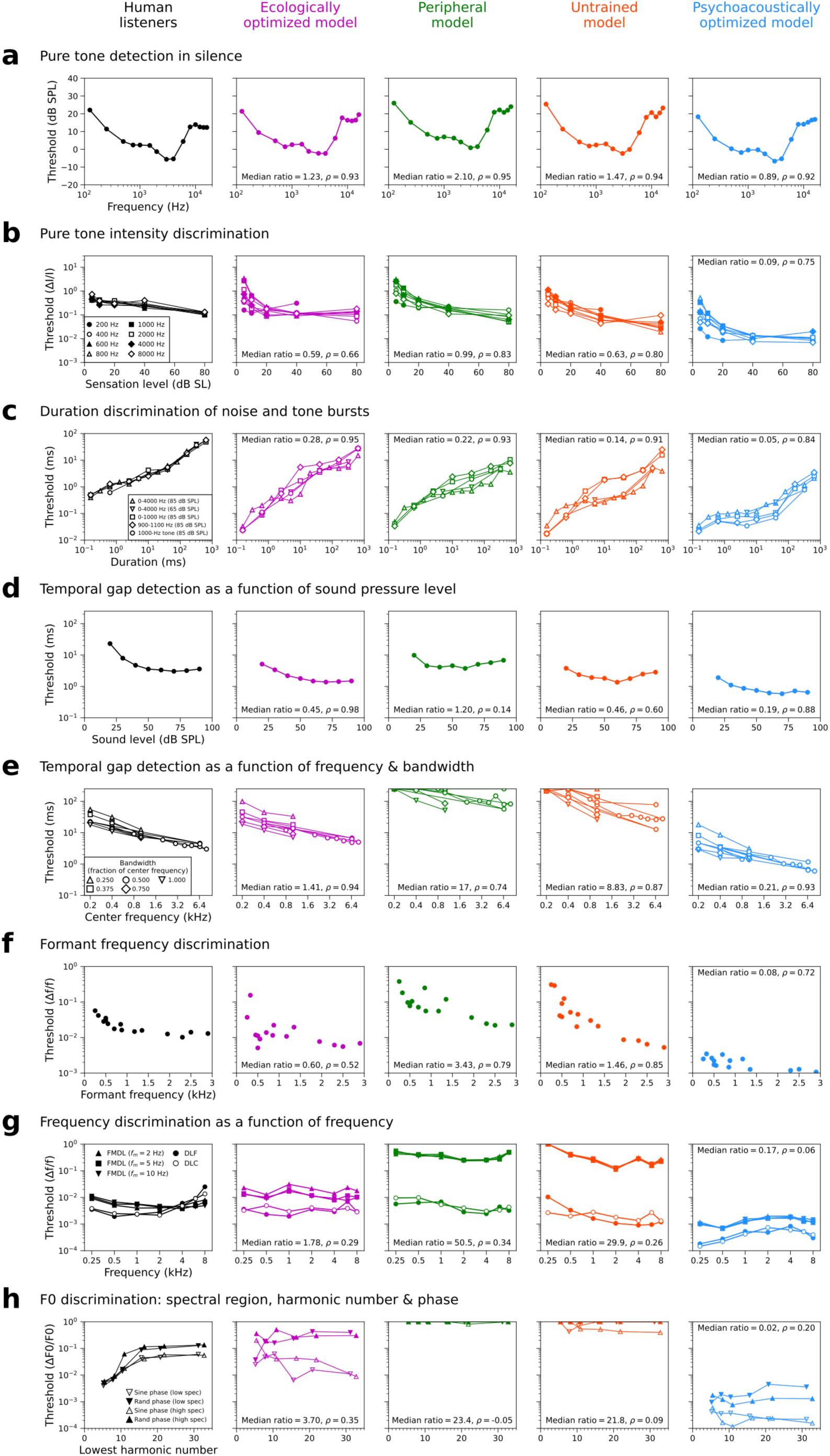
Remaining psychoacoustic experiments. (**a**) Pure tone detection thresholds as a function of frequency. (**b**) Pure tone intensity discrimination thresholds as a function of sensation level. (**c**) Duration discrimination thresholds as a function of stimulus condition and duration. (**d**) Temporal gap detection thresholds for broadband noise as a function of sound level. (**e**) Temporal gap detection thresholds for narrowband noise as a function of center frequency and bandwidth. (**f**) Formant frequency discrimination thresholds for the first and second formants of ten vowels as a function of formant frequency. (**g**) Frequency discrimination thresholds measured as a function of frequency with three different tasks: frequency change detection (DLC), up/down frequency discrimination (DLF), and frequency modulation detection (FMDL). Frequency modulation detection thresholds were measured for three different frequency modulation rates. (**h**) Complex tone F0 discrimination thresholds as a function of spectral region, lowest harmonic number, and harmonic phase. The low (2.5 to 3.5 kHz) and high (5 to 7 kHz) spectral regions had fixed passbands, and the lowest harmonic number was varied by changing F0.

## REFERENCES

1. Campbell, F. W. & Robson, J. G. Application of Fourier analysis to the visibility of gratings. J. Physiol. 197, 551–566 (1968).

2. Hurvich, L. M. & Jameson, D. An opponent-process theory of color vision. Psychol. Rev. 64, 384–404 (1957).

3. Durlach, N. I. Equalization and cancellation theory of binaural masking-level differences. J. Acoust. Soc. Am. 35, 1206–1218 (1963).

4. Dau, T., Kollmeier, B. & Kohlrausch, A. Modeling auditory processing of amplitude modulation. I. Detection and masking with narrow-band carriers. J. Acoust. Soc. Am. 102, 2892–2905 (1997).

5. Barlow, H. B. Retinal noise and absolute threshold. J. Opt. Soc. Am. 46, 634–639 (1956).

6. Green, D. M. & Swets, J. A. Signal Detection Theory and Psychophysics. xi, 455 (John Wiley, Oxford, England, 1966).

7. Faisal, A. A., Selen, L. P. J. & Wolpert, D. M. Noise in the nervous system. Nat. Rev. Neurosci. 9, 292–303 (2008).

8. Barlow, H. B. Possible principles underlying the transformation of sensory messages. Sens. Commun. 1, 217–233 (1961).

9. Attneave, F. Some informational aspects of visual perception. Psychol. Rev. 61, 183– 193 (1954).

10. Olshausen, B. A. & Field, D. J. Emergence of simple-cell receptive field properties by learning a sparse code for natural images. Nature 381, 607–609 (1996).

11. Simoncelli, E. P. & Olshausen, B. A. Natural image statistics and neural representation. Annu. Rev. Neurosci. 24, 1193–1216 (2001).

12. Lewicki, M. S. Efficient coding of natural sounds. Nat. Neurosci. 5, 356–363 (2002).

13. Geisler, W. S. Visual perception and the statistical properties of natural scenes. Annu. Rev. Psychol. 59, 167–192 (2008).

14. Ganguli, D. & Simoncelli, E. P. Efficient sensory encoding and bayesian inference with heterogeneous neural populations. Neural Comput. 26, 2103–2134 (2014).

15. Wei, X.-X. & Stocker, A. A. Lawful relation between perceptual bias and discriminability. Proc. Natl. Acad. Sci. 114, 10244–10249 (2017).

16. Kell, A. J. E., Yamins, D. L. K., Shook, E. N., Norman-Haignere, S. V. & McDermott, J. H. A task-optimized neural network replicates human auditory behavior, predicts brain responses, and reveals a cortical processing hierarchy. Neuron 98, 630–644.e16 (2018).

17. Saddler, M. R. & McDermott, J. H. Models optimized for real-world tasks reveal the task-dependent necessity of precise temporal coding in hearing. Nat. Commun. 15, 10590 (2024).

18. Rajalingham, R. et al. Large-scale, high-resolution comparison of the core visual object recognition behavior of humans, monkeys, and state-of-the-art deep artificial neural networks. J. Neurosci. 38, 7255–7269 (2018).

19. Geirhos, R. et al. Partial success in closing the gap between human and machine vision. in Advances in Neural Information Processing Systems vol. 34 23885–23899 (2021).

20. Saddler, M. R., Gonzalez, R. & McDermott, J. H. Deep neural network models reveal interplay of peripheral coding and stimulus statistics in pitch perception. Nat. Commun. 12, 7278 (2021).

21. Francl, A. & McDermott, J. H. Deep neural network models of sound localization reveal how perception is adapted to real-world environments. Nat. Hum. Behav. 6, 111–133 (2022).

22. Griffith, I. M., Hess, R. P. & McDermott, J. H. Optimized feature gains explain and predict successes and failures of human selective listening. Nat. Hum. Behav. 1–23 (2026) doi:10.1038/s41562-026-02414-7.

23. Govindarajan, L. N., Alavilli, S. & McDermott, J. H. Task-optimized models of sensory uncertainty reproduce human confidence judgments. 2025.10.31.685933 Preprint at 10.1101/2025.10.31.685933 (2026).

24. Benjamin, A. S., Zhang, L.-Q., Qiu, C., Stocker, A. A. & Kording, K. P. Efficient neural codes naturally emerge through gradient descent learning. Nat. Commun. 13, 7972 (2022).

25. Alavilli, S. & McDermott, J. H. From sound to source: human and model recognition of environmental sounds. 2026.03.12.711349 Preprint at 10.64898/2026.03.12.711349 (2026).

26. McPherson, M. J. & McDermott, J. H. Diversity in pitch perception revealed by task dependence. Nat. Hum. Behav. 2, 52–66 (2018).

27. Beutelmann, R. & Brand, T. Prediction of speech intelligibility in spatial noise and reverberation for normal-hearing and hearing-impaired listeners. J. Acoust. Soc. Am. 120, 331–342 (2006).

28. Bronkhorst, A. W. The cocktail party phenomenon: a review of research on speech intelligibility in multiple-talker conditions. Acta Acust. United Acust. 86, 117–128 (2000).

29. Viemeister, N. F. Temporal modulation transfer functions based upon modulation thresholds. J. Acoust. Soc. Am. 66, 1364–1380 (1979).

30. Chi, T., Ru, P. & Shamma, S. A. Multiresolution spectrotemporal analysis of complex sounds. J. Acoust. Soc. Am. 118, 887–906 (2005).

31. Dau, T., Verhey, J. & Kohlrausch, A. Intrinsic envelope fluctuations and modulation-detection thresholds for narrow-band noise carriers. J. Acoust. Soc. Am. 106, 2752– 2760 (1999).

32. Ewert, S. D. & Dau, T. Characterizing frequency selectivity for envelope fluctuations. J. Acoust. Soc. Am. 108, 1181–1196 (2000).

33. Eddins, D. A. & Bero, E. M. Spectral modulation detection as a function of modulation frequency, carrier bandwidth, and carrier frequency region. J. Acoust. Soc. Am. 121, 363–372 (2007).

34. Chi, T., Gao, Y., Guyton, M. C., Ru, P. & Shamma, S. Spectro-temporal modulation transfer functions and speech intelligibility. J. Acoust. Soc. Am. 106, 2719–2732 (1999).

35. Saoji, A. A. & Eddins, D. A. Spectral modulation masking patterns reveal tuning to spectral envelope frequency. J. Acoust. Soc. Am. 122, 1004–1013 (2007).

36. Oetjen, A. & Verhey, J. L. Spectro-temporal modulation masking patterns reveal frequency selectivity. J. Acoust. Soc. Am. 137, 714–723 (2015).

37. Yost, W. A. & Dye, R. H., Jr. Discrimination of interaural differences of level as a function of frequency. J. Acoust. Soc. Am. 83, 1846–1851 (1988).

38. Brughera, A., Dunai, L. & Hartmann, W. M. Human interaural time difference thresholds for sine tones: The high-frequency limit. J. Acoust. Soc. Am. 133, 2839–2855 (2013).

39. Bernstein, L. R. & Trahiotis, C. Enhancing sensitivity to interaural delays at high frequencies by using “transposed stimuli”. J. Acoust. Soc. Am. 112, 1026–1036 (2002).

40. Gabriel, K. J. & Colburn, H. S. Interaural correlation discrimination: I. Bandwidth and level dependence. J. Acoust. Soc. Am. 69, 1394–1401 (1981).

41. Mills, A. W. On the minimum audible angle. J. Acoust. Soc. Am. 30, 237–246 (1958).

42. Hirsh, I. J. The influence of interaural phase on interaural summation and inhibition. J. Acoust. Soc. Am. 20, 536–544 (1948).

43. van de Par, S. & Kohlrausch, A. Dependence of binaural masking level differences on center frequency, masker bandwidth, and interaural parameters. J. Acoust. Soc. Am. 106, 1940–1947 (1999).

44. van de Par, S. & Kohlrausch, A. A new approach to comparing binaural masking level differences at low and high frequencies. J. Acoust. Soc. Am. 101, 1671–1680 (1997).

45. Robinson, D. E. & Jeffress, L. A. Effect of varying the interaural noise correlation on the detectability of tonal signals. J. Acoust. Soc. Am. 35, 1947–1952 (1963).

46. Jesteadt, W., Wier, C. C. & Green, D. M. Intensity discrimination as a function of frequency and sensation level. J. Acoust. Soc. Am. 61, 169–177 (1977).

47. Abel, S. M. Duration discrimination of noise and tone bursts. J. Acoust. Soc. Am. 51, 1219–1223 (1972).

48. Florentine, M. & Buus, S. Temporal gap detection in sensorineural and simulated hearing impairments. J. Speech Lang. Hear. Res. 27, 449–455 (1984).

49. Shailer, M. J. & Moore, B. C. J. Gap detection as a function of frequency, bandwidth, and level. J. Acoust. Soc. Am. 74, 467–473 (1983).

50. Jesteadt, W., Bacon, S. P. & Lehman, J. R. Forward masking as a function of frequency, masker level, and signal delay. J. Acoust. Soc. Am. 71, 950–962 (1982).

51. Oxenham, A. J. & Moore, B. C. J. Modeling the additivity of nonsimultaneous masking. Hear. Res. 80, 105–118 (1994).

52. Jepsen, M. L., Ewert, S. D. & Dau, T. A computational model of human auditory signal processing and perception. J. Acoust. Soc. Am. 124, 422–438 (2008).

53. Greenwood, D. D. Auditory masking and the critical band. J. Acoust. Soc. Am. 33, 484– 502 (1961).

54. Moore, B. C. J., Glasberg, B. R. & Roberts, B. Refining the measurement of psychophysical tuning curves. J. Acoust. Soc. Am. 76, 1057–1066 (1984).

55. Moore, B. C. J., Alcántara, J. I. & Dau, T. Masking patterns for sinusoidal and narrow-band noise maskers. J. Acoust. Soc. Am. 104, 1023–1038 (1998).

56. Kewley-Port, D. & Watson, C. S. Formant-frequency discrimination for isolated English vowels. J. Acoust. Soc. Am. 95, 485–496 (1994).

57. Sek, A. & Moore, B. C. J. Frequency discrimination as a function of frequency, measured in several ways. J. Acoust. Soc. Am. 97, 2479–2486 (1995).

58. Bernstein, J. G. W. & Oxenham, A. J. An autocorrelation model with place dependence to account for the effect of harmonic number on fundamental frequency discrimination. J. Acoust. Soc. Am. 117, 3816–3831 (2005).

59. Oxenham, A. J. & Plack, C. J. Effects of masker frequency and duration in forward masking: further evidence for the influence of peripheral nonlinearity. Hear. Res. 150, 258–266 (2000).

60. Verschooten, E. et al. The upper frequency limit for the use of phase locking to code temporal fine structure in humans: A compilation of viewpoints. Hear. Res. 377, 109– 121 (2019).

61. Houtsma, A. J. M. & Smurzynski, J. Pitch identification and discrimination for complex tones with many harmonics. J. Acoust. Soc. Am. 87, 304–310 (1990).

62. Tuckute, G., Feather, J., Boebinger, D. & McDermott, J. H. Many but not all deep neural network audio models capture brain responses and exhibit correspondence between model stages and brain regions. PLOS Biol. 21, e3002366 (2023).

63. Kubilius, J., Bracci, S. & Beeck, H. P. O. de. Deep neural networks as a computational model for human shape sensitivity. PLOS Comput. Biol. 12, e1004896 (2016).

64. Akbarinia, A., Morgenstern, Y. & Gegenfurtner, K. R. Contrast sensitivity function in deep networks. Neural Netw. 164, 228–244 (2023).

65. Margalit, E. et al. A unifying framework for functional organization in early and higher ventral visual cortex. Neuron 112, 2435–2451.e7 (2024).

66. Grothe, B., Pecka, M. & McAlpine, D. Mechanisms of sound localization in mammals. Physiol. Rev. 90, 983–1012 (2010).

67. Micheyl, C., Delhommeau, K., Perrot, X. & Oxenham, A. J. Influence of musical and psychoacoustical training on pitch discrimination. Hear. Res. 219, 36–47 (2006).

68. Bharadwaj, H. M., Masud, S., Mehraei, G., Verhulst, S. & Shinn-Cunningham, B. G. Individual differences reveal correlates of hidden hearing deficits. J. Neurosci. 35, 2161–2172 (2015).

69. Hung, C. P., Kreiman, G., Poggio, T. & DiCarlo, J. J. Fast readout of object identity from macaque inferior temporal cortex. Science 310, 863–866 (2005).

70. Majaj, N. J., Hong, H., Solomon, E. A. & DiCarlo, J. J. Simple learned weighted sums of inferior temporal neuronal firing rates accurately predict human core object recognition performance. J. Neurosci. 35, 13402–13418 (2015).

71. Hong, H., Yamins, D. L. K., Majaj, N. J. & DiCarlo, J. J. Explicit information for category-orthogonal object properties increases along the ventral stream. Nat. Neurosci. 19, 613–622 (2016).

72. Storrs, K. R., Anderson, B. L. & Fleming, R. W. Unsupervised learning predicts human perception and misperception of gloss. Nat. Hum. Behav. 5, 1402–1417 (2021).

73. Heinz, M. G. Computational modeling of sensorineural hearing loss. in Computational Models of the Auditory System 177–202 (Springer, Boston, MA, 2010). doi:10.1007/978-1-4419-5934-8_7.

74. Brochier, T. et al. From microphone to phoneme: an end-to-end computational neural model for predicting speech perception with cochlear implants. IEEE Trans. Biomed. Eng. 69, 3300–3312 (2022).

75. Saddler, M. R., Dau, T. & McDermott, J. H. Towards individualized models of hearing-impaired speech perception. in Proc. The 6th Clarity Workshop on Improving Speech-in-Noise for Hearing Devices (Clarity-2025) 7–11 (2025). doi:10.21437/Clarity.2025-3.

76. Banerjee, A., Saddler, M. R., Arenberg, J. G. & McDermott, J. H. A deep learning framework for understanding cochlear implants. 2025.07.16.665227 Preprint at 10.1101/2025.07.16.665227 (2025).

77. Newsome, W. T., Britten, K. H. & Movshon, J. A. Neuronal correlates of a perceptual decision. Nature 341, 52–54 (1989).

78. Gold, J. I. & Shadlen, M. N. The neural basis of decision making. Annu. Rev. Neurosci. 30, 535–574 (2007).

79. Arieli, A., Sterkin, A., Grinvald, A. & Aertsen, A. Dynamics of ongoing activity: explanation of the large variability in evoked cortical responses. Science 273, 1868– 1871 (1996).

80. Stringer, C. et al. Spontaneous behaviors drive multidimensional, brainwide activity. Science 364, eaav7893 (2019).

81. Shadlen, M. N. & Newsome, W. T. The variable discharge of cortical neurons: implications for connectivity, computation, and information coding. J. Neurosci. 18, 3870–3896 (1998).

82. Ardila, R. et al. Common Voice: A massively-multilingual speech corpus. in Proceedings of the Twelfth Language Resources and Evaluation Conference (eds Calzolari, N. et al.) 4218–4222 (European Language Resources Association, Marseille, France, 2020).

83. Yadav, S. & Foster, M. E. GISE-51: A scalable isolated sound events dataset. Preprint at 10.48550/arXiv.2103.12306 (2021).

84. Fonseca, E., Favory, X., Pons, J., Font, F. & Serra, X. FSD50K: An open dataset of human-labeled sound events. IEEEACM Trans. Audio Speech Lang. Process. 30, 829–852 (2022).

85. Gemmeke, J. F. et al. Audio Set: An ontology and human-labeled dataset for audio events. in Proc. IEEE ICASSP 2017 (New Orleans, LA, 2017).

86. Gardner, W. G. & Martin, K. D. HRTF measurements of a KEMAR. J. Acoust. Soc. Am. 97, 3907–3908 (1995).

87. Shinn-Cunningham, B. G., Desloge, J. G. & Kopco, N. Empirical and modeled acoustic transfer functions in a simple room: effects of distance and direction. in Proceedings of the 2001 IEEE Workshop on the Applications of Signal Processing to Audio and Acoustics (Cat. No.01TH8575) 183–186 (2001). doi:10.1109/ASPAA.2001.969573.

88. Bruce, I. C., Erfani, Y. & Zilany, M. S. A. A phenomenological model of the synapse between the inner hair cell and auditory nerve: Implications of limited neurotransmitter release sites. Hear. Res. 360, 40–54 (2018).

89. Liberman, M. C. Auditory-nerve response from cats raised in a low-noise chamber. J. Acoust. Soc. Am. 63, 442–455 (1978).

90. Saddler, M. R. et al. Speech denoising with auditory models. in Proc. Interspeech 2021 2681–2685 (ISCA, 2021). doi:10.21437/Interspeech.2021-1973.

91. Köhn, A., Stegen, F. & Baumann, T. Mining the Spoken Wikipedia for speech data and beyond. in Proceedings of the Tenth International Conference on Language Resources and Evaluation (LREC’16) 4644–4647 (European Language Resources Association (ELRA), Portorož, Slovenia, 2016).

92. Woods, K. J. P., Siegel, M. H., Traer, J. & McDermott, J. H. Headphone screening to facilitate web-based auditory experiments. Atten. Percept. Psychophys. 79, 2064–2072 (2017).

93. Kawahara, H. et al. Tandem-STRAIGHT: A temporally stable power spectral representation for periodic signals and applications to interference-free spectrum, F0, and aperiodicity estimation. 2008 IEEE Int. Conf. Acoust. Speech Signal Process. 3933–3936 (2008) doi:10.1109/icassp.2008.4518514.

94. Kollmeier, B. et al. The multilingual matrix test: Principles, applications, and comparison across languages: A review. Int. J. Audiol. 54, 3–16 (2015).

95. ISO 226:2003 Acoustics — Normal equal-loudness-level contours.

96. Resnick, J. M., Horn, D. L., Noble, A. R. & Rubinstein, J. T. Spectral aliasing in an acoustic spectral ripple discrimination task. J. Acoust. Soc. Am. 147, 1054–1058 (2020).

